# Convergent genetic rewiring of the brain underlies termite sociality

**DOI:** 10.64898/2025.12.26.695997

**Authors:** Alina A. Mikhailova, Cédric Aumont, Cong Liu, Aleš Buček, Yi-Ming Weng, Simon Hellemans, Tracy Audisio, Crystal Clitheroe, Ives Haifig, Shulin He, David Sillam-Dussès, Gaku Tokuda, Zongqing Wang, Jan Šobotník, Thomas Bourguignon, Mark C. Harrison, Dino P. McMahon

**Affiliations:** Institute for Evolution and Biodiversity, University of Münster, 48149 Münster, Germany; Department for Materials and the Environment, BAM Federal Institute for Materials Research and Testing, 12205, Berlin, Germany; Institute of Biology, Freie Universitat Berlin, 14195, Berlin, Germany; Okinawa Institute of Science & Technology Graduate University, 904-0495 Okinawa, Japan; Biology Centre, Czech Academy of Sciences, 37005 České Budějovice, Czech Republic; Universidade Federal do ABC, Santo André, SP, 09210-580 Brazil; College of Life Sciences, Chongqing Normal University, Chongqing, People’s Republic of China; University Sorbonne Paris Nord, Laboratory of Experimental and Comparative Ethology UR4443, 93430 Villetaneuse, France; Tropical Biosphere Research Center, University of the Ryukyus, Okinawa, 903-0213, Japan; College of Plant Protection, Southwest University, Chongqing, 400715 China; Faculty of Tropical AgriSciences, Czech University of Life Sciences, 16500 Prague, Czech Republic; Centre for Health & Life Sciences, Coventry University, Coventry, UK

## Abstract

How genomes encode major transitions in social evolution is unclear. We use 29 near-chromosome-quality genomes across a spectrum of social complexity to explore the genomic basis of termite sociality. We show that shifts in selection and gene family evolution preceded the emergence of sociality, pointing to subtler genetic causes of this major evolutionary transition (MET). In comparisons of convergent societal forms, we find that a subset of Odorant Receptors (ORs) underwent parallel expansions in independent advanced termite societies displaying true worker phenotypes. The identified OR genes play caste-differentiated roles in termite but not nearest roach brains and are especially elaborated in true workers. Together with evidence of co-opted nutritional signaling and behavioral genes at different levels of social complexity, our study illuminates the rewiring of the molecular machinery underlying this MET.

---

A major evolutionary transition (MET) entails modifications to the way information is stored and transmitted (Maynard Smith & Szathmáry 1995), resulting in new organisms that possess emergent properties that go beyond the actions of individual components (West et al. 2014, Bar-Yam 2019). Insect societies represent appealing examples of METs due to their formation from relatively simple agents that collectively possess a range of impressive emergent traits (Gordon 2010; Ulrich et al. 2018; Werfel et al. 2014), which in turn explain their ecological dominance in terrestrial ecosystems (Tuma et al. 2019; Ashton et al. 2019; Zanne et al. 2022).

Adaptation towards task specialization via the division of labor represents a major emergent property of insect societies, and among the termites, this has given rise to spectacular examples of polyphenism throughout evolution (Fig. 1a). Owing to their hemimetabolous form of development, termite societies have evolved a diversity of caste phenotypes that differentiate gradually over individual ontogeny (Korb & Hartfelder 2008, Revely et al. 2022). In the most advanced termite societies, bifurcation into dedicated “true” worker (TW) and reproductive phenotypes (Fig. 1b) occurs during development (Revely et al. 2024, Roisin 2000) whereas all other termites possess a linear form of development comprised of totipotent “false” workers (FWs). We performed ancestral state reconstruction of worker type over termite phylogeny using the best fitting tree model (Akaike weight = 0.96) from Hellemans et al. (2024) to understand the evolutionary history of termite workers and to provide a comparative framework for subsequent analyses. Our reconstruction supports a scenario in which TWs evolved four times independently from an ancestrally linear form of development (Fig. 1c, Fig. S1).

**Fig. 1.**
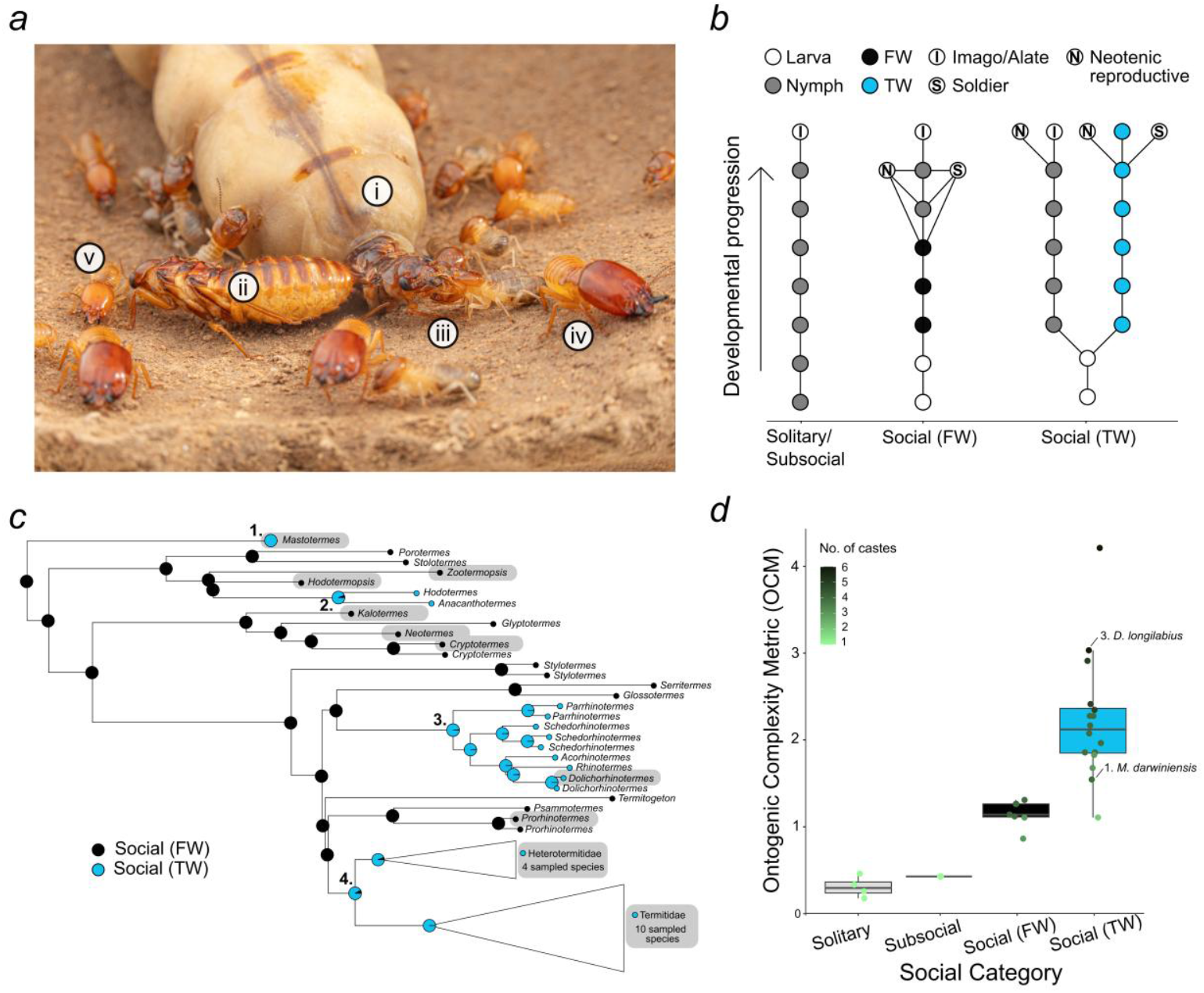
**(a)** Extreme polyphenism in the termite *Macrotermes michaelseni* (Termitidae). The following castes are visible: (i) queen, (ii) king, (iii) workers, (iv) major and (v) minor soldier. **(b)** Simplified ontogeny schemes according to social category with nodes representing instars connected by progressive moults. Solitary and subsocial refer to cockroaches and wood roaches, respectively, while FW and TW phenotypes correspond to linear and bifurcated termite ontogenies, respectively. For simplification, some termite network connections, such as regressive and intermediate moults or caste variants, are not shown. Neotenic reproductives attain reproductive maturity without going through complete somatic (e.g. eye) differentiation and can be derived from either the linear/nymphal or the true worker line. **(c)** Reconstruction of the worker phenotype over termite phylogeny (from Hellemans et al. 2024, Fig. S1) supports four independent origins of true workers (TWs) in termites, labelled 1-4 for each node corresponding to the reconstructed TW transition.Alternative hypotheses proposing TWs as ancestral have previously been considered (Thompson et al. 2000, Bourguignon et al. 2016) but our analysis supports a convergent model of TW evolution (Roisin & Korb 2011). Genera sampled in this study are highlighted in grey. The families Heterotermitidae and Termitidae (collectively termed Geoisoptera) have each been collapsed to ease visualization. **(d)** Significant differences in complexity are discernible between termite social categories, as quantified by the *OCM*, which measures the average number of molts between all instars (*SPL*) corrected by the number of terminal stages. For this calculation, TWs but not FWs were considered a terminal stage. Termite social categories also differed based on the *SPL* metric alone, with TW species harboring a significantly lower *SPL* than FW species (posterior mean = −1.03, 95% CI = [−1.44] − [−0.63]). As in (b) solitary and subsocial categories refer to cockroaches and wood roaches respectively. Social (TW) species belong to TW node 4, except where indicated (node 1: *Mastotermes dawiniensis*, node 3: *Dolichorhinotermes longilabius*).

To quantify the relationship between termite caste differentiation and developmental potency, we developed an Ontogenic Complexity Metric (*OCM*), which measures the shortest characteristic path length (*SPL*) of a species’ developmental network (Fig. 1b), corrected by the number of terminal stages (Supplementary Text), and plotted these against four *a priori* social categories (Fig. 1d). These reveal convergent changes in termite social complexity, with TW species showing a higher *OCM* than FW species, with an MCMCglmm posterior mean of 1.06 and a 95% credible interval (CI) of 0.22 to 1.81, which excludes zero and therefore supports a credible difference (Supplementary Material).

To examine the genetic basis of the transition to termite sociality, we carried out a comparison of near-chromosome-quality genomes from 29 cockroach and termite species spanning a spectrum of social complexity from solitary cockroaches, subsocial wood roaches and a range of social termite species including three independent TW lineages (corresponding to tree nodes 1., 3., and 4., Fig. 1c) (Liu et al. 2025). Caste-specific gene expression patterns from the brains of a subset of 14 representative species were sequenced to investigate the evolution of genetic mechanisms regulating termite caste differentiation.

## Major genomic shifts in selection predate the origin of termite sociality

We found that a major genomic indicator of termite sociality - the relaxation of selection, which may be linked to reduced effective population sizes of social organisms (Ewart et al. 2024) - predated the origin of termites, instead being linked to the emergence of subsocial living in the common ancestor of *Cryptocercus* wood roaches and termites (Fig. 2a-d), confirming a recent finding (Jones et al. 2025). In an analysis of single-copy orthologs (SCOs), median species values of the evolutionary parameter *k*, which measures the degree of intensified (*k*>1) versus relaxed (*k*<1) selection, were above or approximately one for solitary cockroaches (median 1.12, Inter-Quartile Range (IQR) 0.81-2.53), and using MCMCglmms, significantly higher compared to *Cryptocercus* (median 0.88, IQR 0.5-1.35, posterior mean= 0.59, 95% Credible Interval (CI) = [0.08 − 1.12]) as well as FW (median 0.75, IQR 0.47-1.10, posterior mean = 0.78, 95% CI = [0.30 − 1.16]), and TW species (median 0.76, IQR 0.44-1.27, posterior mean = 0.76, 95% CI = [0.36 − 1.19]) (Fig. 2a) (Table S1). Values of *k* did not differ significantly between *Cryptocercus* and either of the termite social categories, nor was it significantly associated with *OCM* when accounting for phylogeny. *Cryptocercus* and termite species displayed more genes under significantly relaxed selection and substantially fewer under intensified selection than solitary cockroaches (Fig. 2b). TW termite species displayed a wide range of median *k* per species (from 0.61 to 0.91) (Fig. 2a), as we also found for values of *OCM* (Fig. 1d), as well as a relatively higher number of genes under positive selection, particularly in species belonging to the Termitidae crown group (Fig. 2b). Ten and five Hierarchical Orthologous Groups (HOGs) convergently underwent relaxed and intensified selection respectively on all three sampled TW transitions. However, subsampling at three random branches revealed these values to be within expected distributions of relaxed (62^nd^ percentile) and intensified (13^th^ percentile) gene frequencies, respectively (Fig. 2c). A gene related to membrane-bound transcription factor site-2 protease (S2P), which cleaves transcription factors involved in fatty acid metabolism (Rawson 2003), underwent positive selection at all three TW transitions, suggesting a possible role for fatty acid metabolism in caste-differentiated chemical communication and reproductive traits of TW species.

**Fig. 2.**
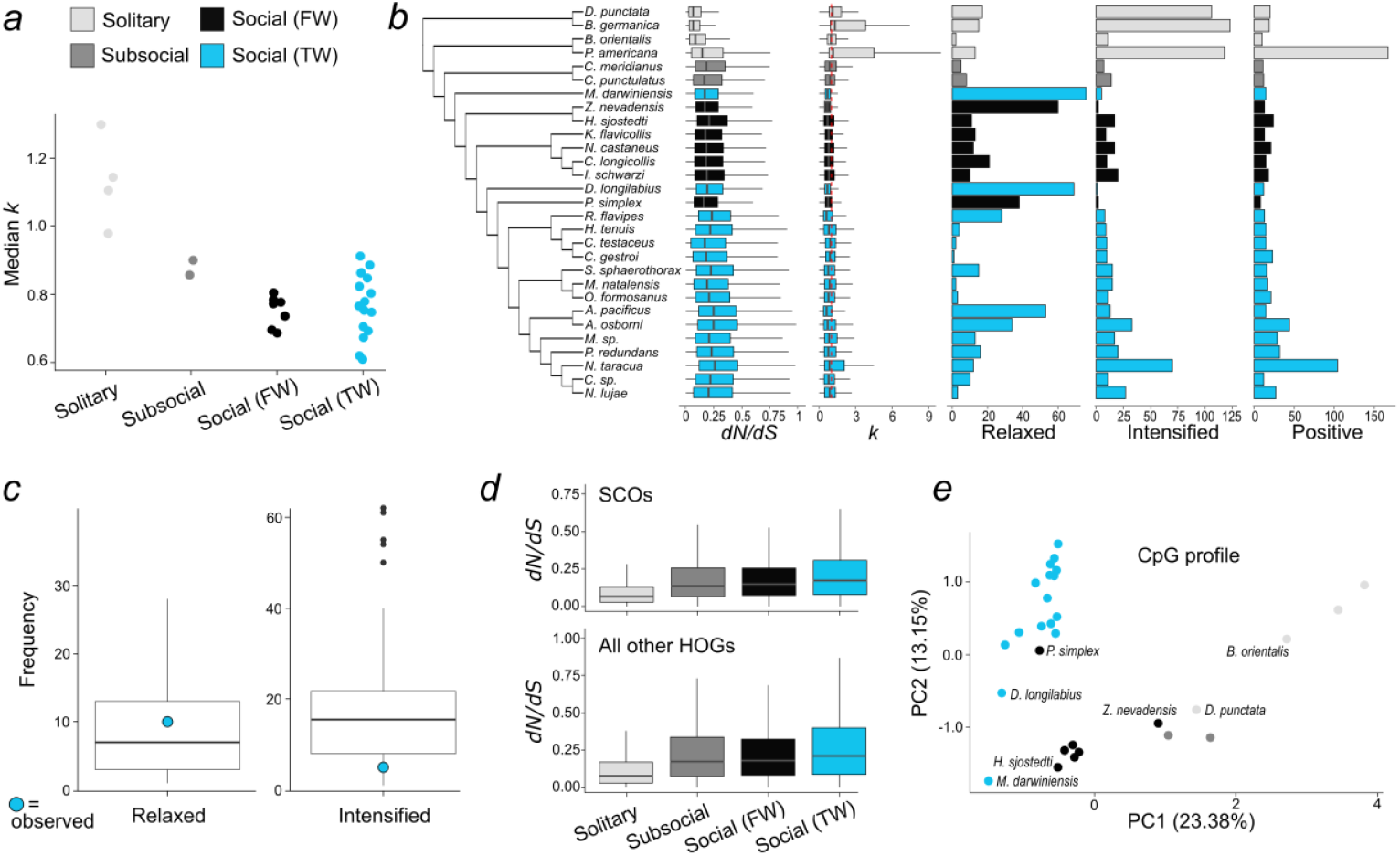
Patterns of selection across the termite MET. **(a)** Median species values of *k* by social category. **(b)** Per species values plotted against a cladogram of the sampled genomes of (left to right): *dN/dS, k*, number of genes identified under relaxed, intensified, and positive selection. For *dN/dS* and *k*, boxplots represent the first and third quartiles as lower and upper hinges, respectively, and 1.5*IQR as whiskers, with the red line (*k*) indicating a value of 1. **(c)** Distributions of relaxed and intensified gene frequencies of 3 randomly selected branches with observed values for the 3 TW transitions (tree nodes 1., 3. and 4. in Fig. 1c) shown in blue. **(d)** *dN/dS* values by social category for SCOs (upper graph) and all other HOGs (lower graph). Boxplots are as in panel (a). **(e)** Principal component analysis of CpG depletion patterns (observed versus expected number of CpGs: CpGo/e) by social category.

Estimated *dN/dS* ratios supported these broad patterns (Fig. 2d, Table S1) with MCMCglmms of SCOs for solitary cockroaches (median *dN/dS* = 0.06, IQR 0.03-0.13) being significantly lower than TW termite species (median *dN/dS* = 0.17, IQR 0.08-0.31, posterior mean = −0.65, 95% CI = [−1.31] − [−0.10]). Estimated *dN/dS* ratios for both *Cryptocercus* and FW termite species overlapped largely with TW termite species and were higher than solitary cockroaches, although not significantly so (Table S1). A similar pattern was found for *dN/dS* of remaining HOGs (excluding SCOs) (Fig. 2d, Table S1).

Overall, these data demonstrate that a significant selective shift occurred in the common ancestral genome of *Cryptocercus* and termites and was not associated with sociality *per se*, with some evidence of a wider range of selective pressures acting on genes in termites with increased social complexity and ecological diversity. DNA methylation patterns from SCOs, as estimated by CpG depletion patterns (observed versus expected number of CpGs: CpGo/e) clustered broadly by social category as well as phylogenetic distance, indicating that social transitions were accompanied by shifts in transcriptional regulation.

## Parallel expansions of chemical communication genes in convergent social transitions

We used CAFE5 (best fitting model, λ = 0.24 with three categories of γ) to analyze gene family expansions and contractions. We identified a trend of gene family contractions across deep *Cryptocercus* + termite branches, in line with their reduced genome size compared with solitary cockroaches, but otherwise detected limited evidence for gene family contraction or expansion at the major social transitions. Instead, we found significantly expanded gene families at three consecutive ancestral nodes of Geoisoptera (N = 28, 3.16 per MY),

Termitidae (N = 52, 3.08 per MY) and Termitidae excluding (Macrotermitinae + Sphaerotermitinae) (N = 50, 8.57 per MY) (Fig. S2). Geoisoptera represents the crown-group of termites that putatively evolved TW once at its common ancestor (node 4 in Fig. 1c). The Termitidae represents the major radiation of termites (Hellemans et al. 2024) and is defined by key shifts in termite symbiosis and diet (Aanen & Eggleton 2017, Chouvenc et al. 2021). The important sub-node comprising all Termitidae except (Macrotermitinae + Sphaerotermitinae) is linked to the evolution of soil feeding (Buček et al. 2019) and is associated with the highest number of gene family expansions, including carbohydrate-active enzymes. This is consistent with general increases in gene number as well as genome size in the most ecologically diverse and species rich termite crown-group (Liu et al. 2025).

To understand if any gene families were specifically linked to convergent increases in social complexity, we carried out a series of phylogenetic models (MCMCglmm) of 569 significantly expanded or contracted gene families. A range of models were used to test the effect of ontogeny or social category on gene family evolution (Supplementary methods). We identified 44 significant gene families overall, of which 10 were shared by representatives from all three transitions to the TW phenotype (Fig. S3, Table S2, Supplementary methods) (N=5 expansions, N=5 contractions). Of these, expansions of two HOGs were supported across all phylogenetic models (Fig. S3). One was identified as a family of G-protein coupled receptors (GPCRs) with sequence similarity to receptors of brain-gut neuropeptides (CCHamide, Ida et al. 2012) (Fig. S4), suggesting co-option of important nutritional or growth-related signalling machinery (Kay et al. 2025) associated with TW species evolution (Fig. S1).

The second was a set of Odorant Receptor (OR) genes which have been implicated in chemical communication innovations in one other social insect MET: the ants (Vizueta et al. 2025, Pellen et al. 2025). Re-annotation and characterization of all ORs by Orthogroup (OG) assignment using Possvm over an OR gene tree revealed two termite-specific OR expansions comprised of a single OR-OG (OR-OG112; OR-cluster 1, Fig. 3a) and a second expansion comprised of multiple OR-OGs (OR-cluster 2, Fig. 3a). Phylogenetic models identified significant enrichment of OR-cluster 1 in all three TW transitions compared with FW species (mean number of ORs per TW species = 20.94 and FW species = 10.43, posterior mean = −10.09, 95% CI = [−17.22] − [−2.55]) (Fig. 3b, Table S3). An expansion of *Mastotermes darwiniensis* (Fig. 1c. TW node 1) OR genes within cluster 1 is especially notable (32 ORs). OR-cluster 2 was also significantly expanded in TW versus FW species, including expansions in TW nodes 3 and 4 (Fig. 1c), but not in *M. darwiniensis* (mean number of ORs per TW species = 25.88 and FW species = 3.00, *M. darwiniensis* = 3, posterior mean = −17.90, 95% CI = [−28.93] − [−6.95]) (Fig. 3b, Table S3). Two further small OR-OGs were significantly expanded in TW termites as were total ORs (mean number of ORs per TW species = 104.69 and FW species = 72.71, posterior mean = −29.66, 95% CI = [−49.33] − [−8.46]) (Fig. 3b, Table S3).

**Fig. 3.**
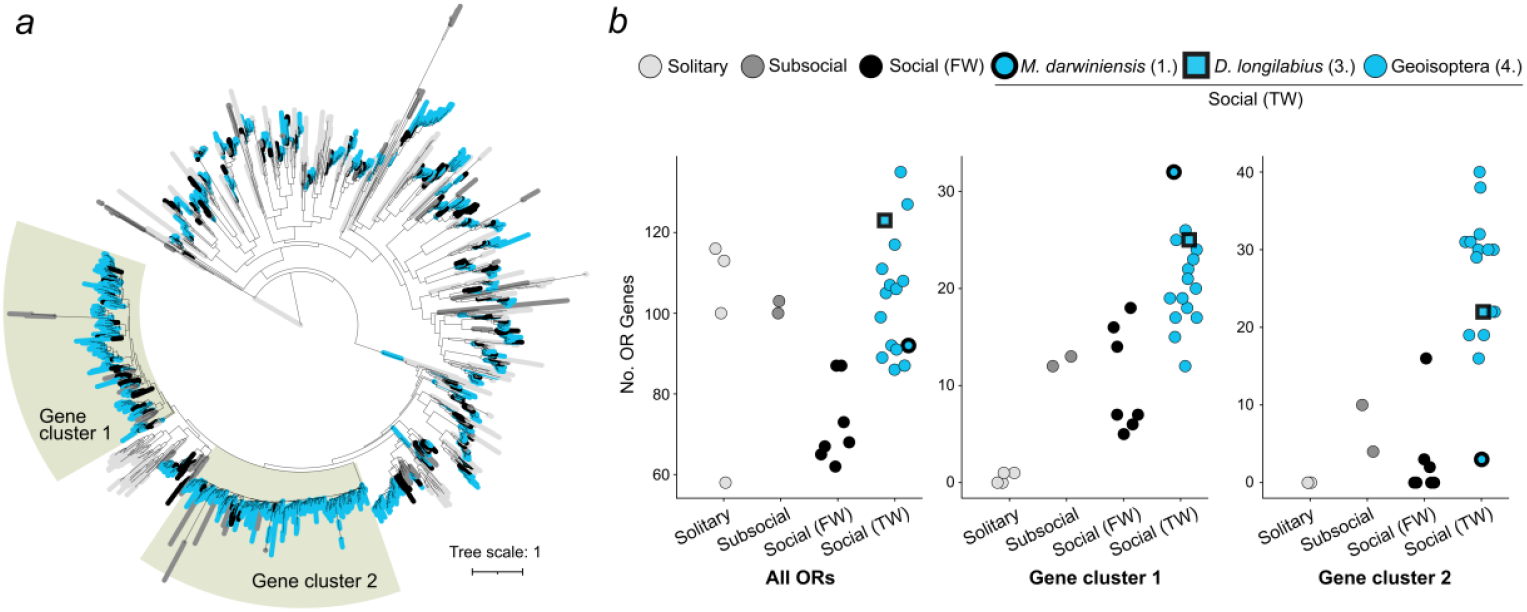
Expansions of Odorant Receptor (OR) genes in termite lineages that evolved the TW phenotype. **(a)** Gene tree of total annotated OR genes across sampled genomes, with 2 major gene expansions (gene cluster 1 and 2) highlighted. Terminal branches are color coded by social category, as in Fig. 2. **(b)** Total OR genes (left panel) are significantly expanded in TW versus FW termites, driven by major expansions in gene cluster 1 (middle panel) and gene cluster 2 (right panel). Species representing independent transitions to the TW phenotype (numbered according to Fig. 1c) are highlighted: Bold circle = *M. darwiniensis*. Bold square = *D. longilabius*. Normal circle = Geoisoptera. *M. darwiniensis* is highly expanded in gene cluster 1 but depauperate (for its social category) in gene cluster 2.

Notably, all but one solitary roach displayed a comparable number of total ORs to subsocial roaches and TW termites yet all are depauperate for OR-cluster 1 and 2, whereas subsocial wood roaches harbored a similar repertoire to FW termites species. Selection analyses of ORs further revealed that multiple ORs from gene clusters 1 and 2 showed signatures of positive selection, but exclusively in TW species (7 and 3, respectively), including three *M. darwiniensis* ORs from OR-cluster 1 (Table S4).

These findings argue that elaborations to specific gene families rather than large-scale shifts in genome architecture or selection are a prevailing feature of termite social evolution.

## Conserved rewiring of gene co-expression architecture linked to chemical communication

Termite castes display significant differences in brain anatomy linked to functional specialization and division of labor (O’Donnell et al. 2022, Ishibashi et al. 2023, Merchant & Zhou 2024). We therefore next compared brain gene co-expression networks to identify signatures of genetic re-wiring linked to changes in cognition or communication during the termite MET. In an initial step, Gene Ontology term analyses showed “sensory perception of smell” (GO:0007608) to be the only significantly (p < 0.01) enriched GO term in caste-specific networks for all but one termite species (8/9 termite species). This enrichment was attributable to the presence of multiple caste-specific ORs. In pairwise comparisons of worker- and reproductive-specific networks, along with a zinc finger protein, we found OR-cluster 1 in all sampled termite species (Super Exact Test (SET): worker network: N=9, F = 252.10, p < 0.001, reproductive network: N=9, F= 251.73, p < 0.01). By contrast, in cockroach network comparisons, OR-cluster 1 was only detected in adult wood roach networks, and the overlap was not significant (SET: F= 0.99, p = 0.70). Similarly, OR-cluster 2 was shared in all reproductive networks of TW species (N=5, SET: F = 5.85, p < 0.001) and in worker networks of 4/5 TW species, not including *M. darwiniensis* (F= 1.85, p < 0.001). In the FW species on the other hand, OR-cluster 2 was only found in 1/4 and 2/4 reproductive and worker networks, respectively. Among the cockroaches, OR-cluster 2 was only shared in the juvenile networks of wood roaches, and again, not significantly so (N=2, SET: F = 1.01, p = 0.35). The reduced representation of OR-cluster 2 in *M. darwiniensis* and FW termite networks reflects the lack of expansion of these genes in these species’ genomes, indicating a specific evolutionary association between OR-cluster 2 and more derived termite TW transitions (nodes 3. and 4., Fig. 1c, Fig. 3b). In SET comparisons including soldiers (Supplementary Methods), we further found that OR-cluster 1 was significantly shared by most termite castes, including soldiers (false workers: 4/5, true workers: 5/5, soldiers: 8/9, primary reproductives: 5/6, and secondary reproductives: 4/5), supporting a central role for OR-cluster 1 in ancestral termite caste-determination.

Finally, comparing OR activity in worker and reproductive co-expression networks across species (Fig. 4a, Fig. S5), we found that the number of ORs in worker networks was greater in 4/5, 3/5 and 3/5 TW versus FW species for total ORs, OR-cluster 1 and 2, respectively, with *M. darwiniensis* being highly represented in total ORs and OR-cluster 1 (Fig. 4b). In an analysis of connectivity between ORs and differentially expressed genes (DEGs) (see below), the TW species *Anoplotermes pacificus, Coptotermes gestroi, Reticulitermes flavipes* and *Macrotermes natalensis* showed the highest degree of connectivitiy between ORs and shared DEGs for total ORs, OR-cluster 1 and 2 (mean connectivity = 64%, 56% and 63%, respectively) (Table S5). Patterns of OR activity and DEG connectivity were less pronounced in corresponding reproductive networks, aside from OR-cluster 2, which was more highly represented and more strongly connected to DEGs in TW species except *M. darwiniensis* (Fig. 4b).

**Fig. 4.**
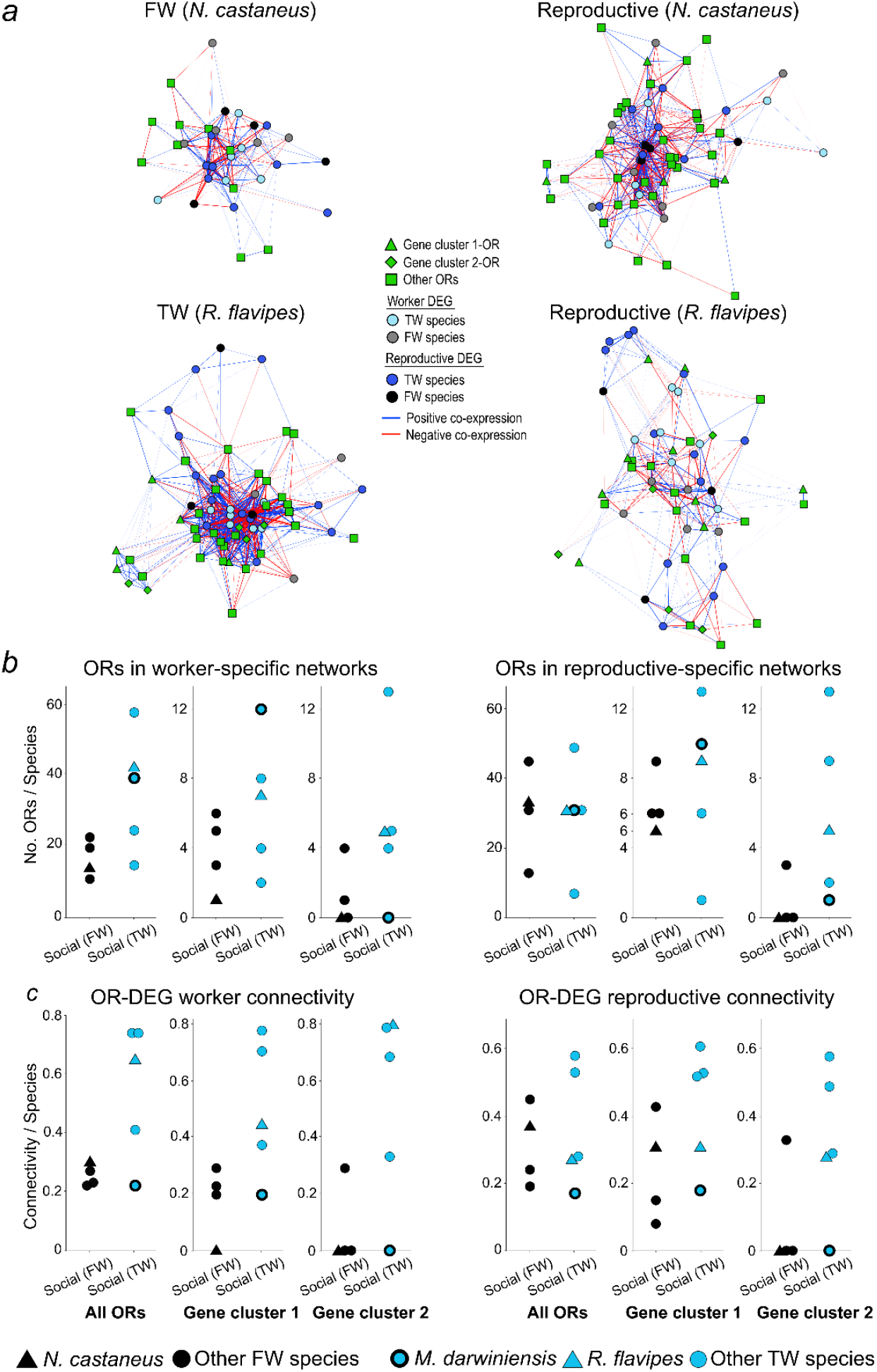
OR recruitment to termite brain gene co-expression networks. **(a)** Simplified caste-specific gene co-expression networks from an FW (*Neotermes castaneus*) and TW (*Reticulitermes flavipes*) termite species highlighting ORs and their connectivity to shared caste-biased DEGs. Left: worker networks. Right: reproductive networks. Nodes represent individual genes. Green: ORs (triangle = gene cluster 1, diamond = gene cluster 2, square = other ORs). Light blue: TW-biased DEG, Grey: FW-biased DEG, Dark blue: reproductive-biased (TW species) DEG, Black: reproductive-biased (FW species) DEG. Edges indicate significant co-expression, with thickness and color representing strength (weight) and direction of co-expression, respectively, Red: negative. Blue: positive. Only edge weights ≥ 0.4 and nodes with at least one apparent connection are represented. **(b)** Number of ORs involved in worker-(left panel) or reproductive-specific (right panel) networks by social category (FW or TW) and OR category. Color coding as in Fig. 3. **(c)**Degree of connectivity between caste-specific ORs and DEGs in worker-(left panel) or reproductive-specific (right panel) networks by social category (FW or TW) and OR category. *M. darwiniensis* node connectivity (22%, 19% and 0%) is closer to species with linear ontogeny for total ORs, OR-cluster 1 and 2 (mean = 26%, 17% and 7%, respectively) (Table S5).

Together with the expansion of ORs, and in particular OR-cluster 1, across three independent termite lineages that evolved the TW phenotype (Fig. 3b), these observed brain gene co-expression patterns support a stepwise model of OR-mediated information signaling evolution in the termite MET: 1) expansion of OR-cluster 1 and 2 in the common ancestor of subsocial roaches and termites, 2) ancestral recruitment of OR-cluster 1 across castes in termite brain evolution, 3) bouts of both parallel and lineage-specific OR gene expansion in TW species, with OR-cluster 2 being specifically linked with derived TW transitions, 4) integration of expanded ORs from both gene clusters into the brain activities of TWs, which are strongly connected to caste-differentiated DEGs.

This scenario is consistent with known anatomical specializations in both the antennal lobes and mushroom bodies of termite workers (O’Donnell et al. 2022, Merchant & Zhou 2024), which receive input from olfactory sensory neurons and act as centers for learning and memory, respectively (Hansson & Stensmyr 2011). Related brain regions are enlarged in the worker castes of divergent hymenopteran societies (Groh & Rössler 2008, Kuebler et al. 2010). In ants, this is also correlated with more extensive OR gene expansion and elaboration (Zhou et al. 2015), with most species harboring several hundred ORs (Pellen et al. 2025), constituting possible evidence of a convergent mechanism of social cognitive specialization over deep evolutionary time.

## Conservation and co-option in the social termite molecular toolkit

Molecular toolkits are evolutionarily conserved genes that are co-opted from solitary ancestors to regulate social insect polyphenism via differential expression or timing between castes (Kocher & Kingwell 2024). We carried out a comparative analysis of brain gene expression levels to identify conserved genes involved in the regulation of termite castes as well as search for evidence of gene co-option under a convergent model of TW evolution. For each species, we idenfied DEGs through pairwise comparisons between worker, soldier and reproductive castes. Inspection of normalized expression patterns of these DEGs across species showed clustering based on caste rather than species, with both FWs and TWs aligning more closely with soldiers than with reproductive castes, regardless of reproductive type, with a weaker effect of sex across caste (Fig. 5a, Fig. S6). In the following section, we concentrate on the identity and putative functions of DEGs in comparisons between workers and reproductives. Selection (*dN/dS* ratios) and Transposable Element (TE) content in proximal (upstream) regulatory regions of DEGs between workers and reproductives were also explored and are reported in Fig. S7 and Table S6.

**Fig. 5.**
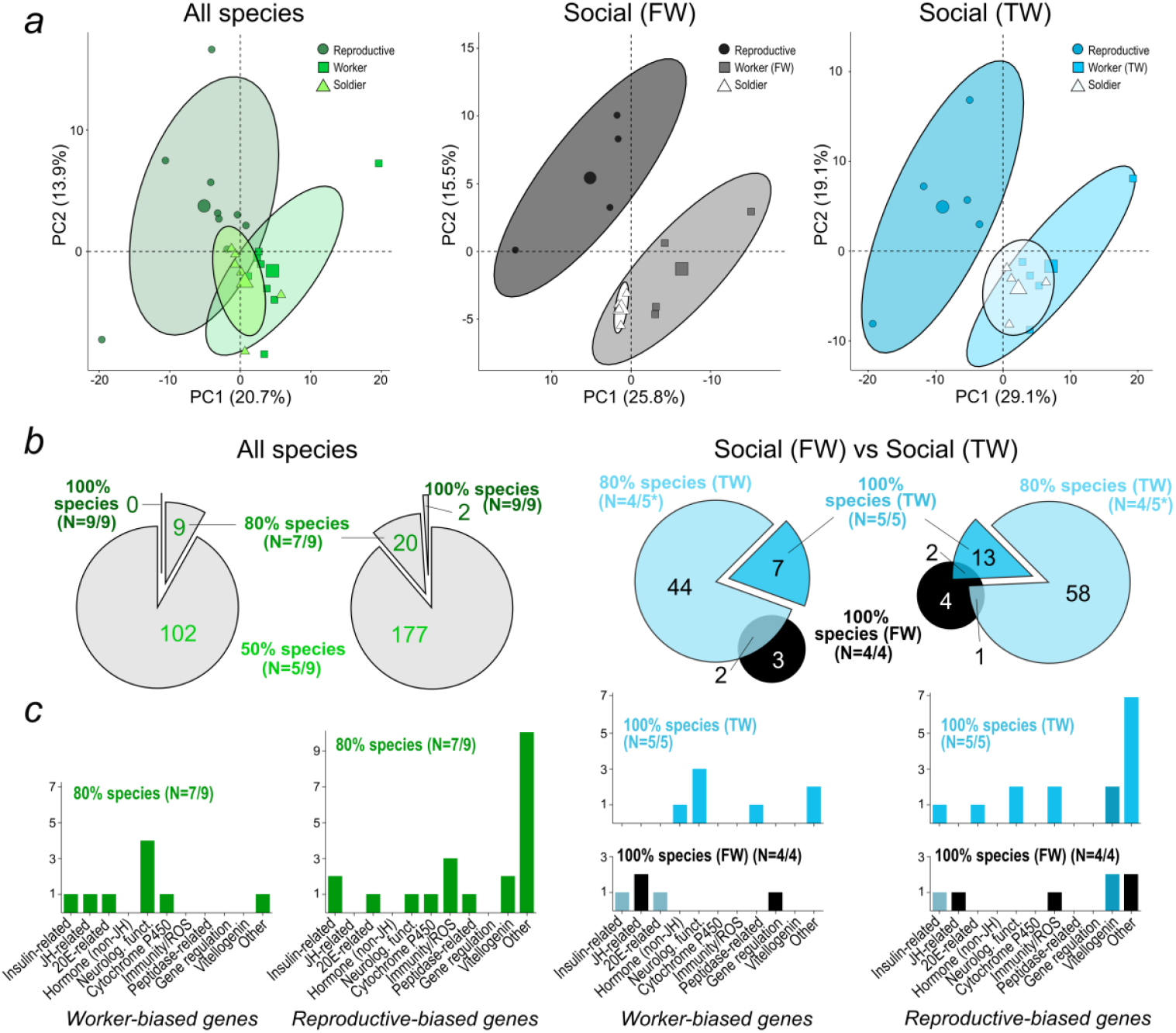
Patterns of gene expression in the termite brain. **(a)** Caste-differentiated PCAs of overexpressed-single copy orthologs in 9 or fewer termite species in: all termite species (left panel), termite species with linear (FW) (middle panel) and bifurcated (TW) development (right panel). **(b)** Shared significantly biased genes in pairwise comparisons of termite workers and reproductive for: all species (left panels) and FW/TW species (right panels). Pie graphs indicate the number of significantly worker- or reproductive-biased genes shared in subsets of all sampled species (100%, 80%, 50% of N=9 species) (left) or specifically FW (100% of N=4) or TW (100%, 80% of N=5) species (right). * DEGs shared by 4/5 TW species include *M. darwiniensis*. **(c)** Classification (see Supplementary Methods) of putative functions of worker- and reproductive-biased genes shared by 80% of all termite species (N=7 of 9) (left panels) and 100% FW (N=4) or TW species (N=5) (right panels). Intermediate colored bars reflect the genes that are overlapping between FW and TW species, as indicated in the pie graphs of (b).

Significantly overlapping sets of orthologs containing DEGs shared by castes across a majority of species were found (Fig. 5b, 5c). Notable among these were 9 HOGs upregulated in both FWs and TWs compared to reproductives in 7/9 species (SET: F > 420, p < 0.003). These 9 HOGs included putative IIS-JH-Vg (insulin-like/IGF-1 signaling (IIS), juvenile hormone (JH), yolk precursor vitellogenin (Vg)) signaling network genes: an Insulin-Like Peptide (ILP), a JH-binding protein, and an Ecdysone-induced protein (Eip74). A Cytochrome P450 (Cyp6a14) and an odor-response related transcriptional regulator (pdm3, Tichy et al 2008) were also identified (Fig. 5c, left).

Of the shared FW and TW-related genes, only 1 HOG (Ecdysone-induced protein) was upregulated in juveniles of solitary cockroaches but not subsocial wood roaches. The near-absence of these genes within lists of roach DEGs points to their possible ancestral recruitment to termite worker regulation, and highlights the importance of rewiring in the IIS-JH-Vg network in particular (Hellemans & Hanus 2024). Inkeeping with these findings, we found two Vitellogenin (Vg) HOGs to be significantly upregulated in reproductives of all termite species (9/9 species comparisons, SET: F = 750278, p < 0.001), while a further 20 were upregulated in reproductives of 7/9 termite species (Fig. 5b, SET: F > 694, p < 0.001).

These included genes related to insulin signaling (Insulin-like peptide, Activin-ß) and immunity/ROS (virus-induced RNA 1, Serine protease, Thioredoxin) (Fig. 5c). Of the two Vg HOGs, one was differentially expressed in adults of at least one solitary cockroach but not in either *Cryptocercus* species. Among the remaining 20 genes, none were upregulated in one or more subsocial or solitary cockroach, further pointing to a conserved role for IIS-JH-Vg network as well as immune signalling associated with termite reproductive function.

Concentrating on TW and FW species separately, we found 7 HOGs to be upregulated in all workers of TW species, including *M. darwiniensis* (N=5 species) (SET: F = 200, p < 0.001). Of these, only a single HOG was upregulated in more than one FW species, indicating a specific association of these genes with the TW phenotype (Table S7). The seven identified HOGs relate to functions in hormonal and behavioural regulation, including putative genes associated with neuronal development (Tomosyn and Src64B), and behaviour (Ktl) (Fig. 5b).

With regard to the FW-related HOGs, while one of the five genes (a transcription factor involved in the development of the central nervous system) was exclusively upregulated in FW species, the remaining four were shared with most TW species and involved in the IIS-JH-Vg pathway, suggestive of a non-specific or ancestral association between these genes and the FW phenotype (Table S7).

Seven and 15 HOGs were shared by reproductives of all FW (SET: F = 25.33, p < 0.001) and TW species (SET: F = 138.60, p < 0.001), respectively. In FW species, these included genes relating to insulin-(Activin-ß) and JH-signaling (hydroxymethylglutaryl-CoA synthase), immunity/ROS (Serine protease), and the 2 Vg HOGs, as described above, while in TW species, immune-(virus-induced RNA 1, Thioredoxin), neuron-related (Moesin, Anosmin-1) and IIS-JH-Vg signaling genes (ILP, an E20-related zinc-finger protein, 2 Vg HOGs) were identified. Unlike worker comparisons, a majority of HOGs associated with reproductives of TW species were shared with more than one FW species (Table S7). This mirrors the unique patterns of OR gene co-expression activity in the workers of TW species, reinforcing the view that specific genetic signatures were associated with TW evolution.

## Conclusions

The evolution of termite sociality was associated with subtle changes to specific gene families and regulatory architecture rather than large scale genomic transformation.

Specifically, our data support a conserved rewiring of IIS-JH-Vg network signalling in early termite evolution, with expansions to the same group of GPCR genes across three independent transitions to TWs pointing to parallel modifications to nutritional signalling in more advanced social lineages. We further show that a subset of OR genes are involved in the regulatory gene co-expression networks of termite brains in a caste-specific manner. Our data indicate that the identified ORs underwent convergent expansions across independent TW termite lineages and that they contribute significantly to the brain gene activity of these species. Alongside evidence of co-opted DEGs linking neuronal development and behavior with the TW phenotype, these findings point to repeated reconfiguration of gene regulatory machinery linked to the specialized sensory and communication functions of true workers.

Comparative functional analysis will be central to efforts seeking to advance understanding of caste cognition as well as nutritional signalling evolution in termites while comparisons with other insect societies will permit exploration of wider toolkit generalities and distinctions between the social insect METs.

## Acknowledgments

We thank Juliette Berger, Sabine Busweiler, Yvonne de Laval, Luiza Helena Bueno da Silva, Ana Maria Costa Leonardo, Anna Prokhorova, Vasiliki Prosgoliti, Alicia Rijlaarsdam and Menglin Wang for rearing and sampling assistance; Katja Nowick for gene co-expression network guidance; Jens Rolff and Robert Paxton for comments on an earlier version of the manuscript. We also thank the DRESDEN-concept Genome Center, supported by the DFG Research Infrastructure Program (Project 407482635) and part of the Next Generation Sequencing Competence Network NGS-CN (Project 423957469) for NGS data production. The authors also thank the HPC Service of FUB-IT (Bennett et al 2020), Freie Universität Berlin, and HPC Service of PALMA, University of Münster, for computing time as well as OIST’s DNA Sequencing Section (SQC) and the Scientific Computation and Data Analysis Section (SCDA) for assistance with sequencing and providing access to the OIST computing cluster, respectively.

## Author contributions

Conceptualization: DPM, MCH, TB

Methodology: AAM, CA, CL, TA, SH, YMW, SH, CC, AB, JS, DPM, MCH,

TB Analysis: AAM, CA, CL, YMW, AB, MCH

Visualization: DPM, MCH, AAM, CA

Funding acquisition: DPM, MCH, TB, JS

Writing – original draft: DPM, MCH, AAM, CA

Writing – review & editing: All authors

## Supplementary Materials

Materials and Methods

Supplementary Text

Figs. S1 to S7

Tables S1 to S8

References

