## Supplementary Materials for "Convergent genetic rewiring of the brain underlies termite sociality"

**The PDF file includes:**

Materials and Methods
Supplementary Text
Figs. S1 to S7
Tables S1 to S8
References

### Materials and Methods

#### Ancestral state reconstruction

Two distinct types of worker caste have long been recognized in termites, but their origin has been disputed (Thompson et al. 2000; Grandcolas and D'Haese 2002, 2004; Legendre et al. 2013). Using a recent termite phylogeny (Hellemans et al. 2024) we aimed to provide an up-to-date assessment of the ancestral state of termite workers to provide a framework to study the evolution of termite social complexity. The ancestral state of the termite worker caste was reconstructed by considering the worker caste as a discrete binary trait with mutually exclusive states: "true worker" and "pseudergate" (otherwise termed "false worker") (Roisin and Korb 2010). A majority rule consensus of maximum likelihood ultra-conserved element-based trees was used as the phylogenetic hypothesis (tree file "73_iqtree_majority_rule_phytools_updated.tree" available as supplementary in Hellemans et al. (2024)). Zero-length branches corresponding to polytomies were arbitrarily resolved by short internal branches with 1/1000000 of the maximum tree height, i.e. the maximum sum of edge length connecting any tip to the root. The ancestral states were reconstructed using maximum-likelihood methods modelling the evolution of termite worker caste using a time-continuous Markov model (Pagel 1994) as implemented in the "ace" function of the R-package "ape" v5.6-1 (Paradis and Schliep 2019). We used both the model ER (equal rates of evolutionary transitions from pseudergates to true workers and from true workers to pseudergates) and ARD (different rates for the two possible transitions). We also used an ARD model with two and three, respectively, rate categories that allow for different transition rate classes on different portions of a phylogeny. We used this model as implemented in "corHMM" function of the R-package "corHMM" v2.8 (Boyko and Beaulieu 2021). The best-fitting model was selected based on the highest Akaike weight (Wagenmakers and Farrell 2004) which was calculated for the set of all four tested models using the "aic.w" function from the R-package "phytools" v1.2-0 (Revell 2012).

#### Time-calibrated phylogeny

Using the genomes of three solitary cockroaches *B. germanica*, *P. americana*, and *D. punctata* (Harrison et al. 2018; Li et al. 2018; Fouks et al. 2023) and new high-resolution genomes of the solitary cockroach *B. orientalis*, 2 sub-social *Cryptocercus* cockroaches and 23 termites (Jones et al. 2025; Liu et al. 2025) we reconstructed the timescale of termite evolution for use in downstream analysis with the program MCMCTREE, part of the PAML 4.9g package (Yang, 2007). MCMCTREE uses Bayesian estimation of species divergence time with approximate likelihood calculation, which facilitates reconstruction of time-calibrated phylogenies with large numbers of genetic loci. A concatenated protein supermatrix was used as the input sequence alignment. The species tree topology was constructed using IQ-TREE v2.1.3 based on 826 single-copy orthologs with the MFP+MERGE algorithm (Minh et al. 2020). The sequences were aligned with PRANK v.170427 with the option "-F" and concatenated with FASconCAT-G v1.05.1 (Löytynoja 2013, Kück and Longo 2014).

Age calibrations were implemented as lower and upper soft bounds for fourteen internal tree nodes based on termite and cockroach fossils (Table S8). The probability of the calibrated node age being outside of the defined range was set to 2.5%. An additional prior for maximal root age was specified within the MCMCTREE configuration file as 323.2 million years, based on the age of *Mylacris anthracophila*, the first roachoid fossil and stem Dictyoptera (Durden, 1969). Ages of all fossils were retrieved from Paleobiology Database (https://paleobiodb.org/, accessed October 2023). In MCMCTREE, we first calculated the gradient and Hessian of the log-likelihood (option "usedata = 3") with a simple protein model (Poisson without gamma rates). Next, we used empirical rate matrix (WAG) with gamma rates among sites (arguments fix_alpha = 0, alpha = 0.5, ncatG = 4) to generate a new Hessian matrix using this WAG+Gamma model. The matrix values were used for MCMC sampling of posterior distribution using the approximate likelihood method (option "usedata = 2"). We used an independent lognormal clock mode. The MCMC chain was run for 100,000,000 iterations and sampled every 100 iterations. The first 10% iterations were discarded as burn-in. We ran three replicates of the MCMC chain and used Tracer v1.7.1 (Rambaut et al., 2018) to confirm that each replicate had reached sufficient effective sample size and converged to near-completely overlapping node age interval estimates.

**Selection analyses**

Sociality has been hypothesized to be linked to relaxed selection on pleiotropic genes involved in social function (Gadagkar 1997), but more generally, neutral processes such as reductions in effective population size in social organisms have been proposed to influence the strength of purifying selection. This is supported by evidence for weakened purifying selection in social hymenopteran species (Mikhailova et al. 2024) as well as termites (Ewart et al. 2024). We expanded the investigation into selection by exploring the history of selective changes in the genome over cockroach and termite phylogeny, as well as asking how selective forces may be correlated with different levels of termite social complexity, such as the repeated emergence of TW lineages. To carry out selection analysis, we first retrieved the longest isoform from the previously published proteomes of three solitary cockroaches *B. germanica*, *P. americana*, and *D. punctata* (Harrison et al. 2018; Li et al. 2018; Fouks et al. 2023) and new high-resolution genomes of the solitary cockroach *B. orientalis*, two sub-social *Cryptocercus* cockroaches and 23 termites (Jones et al. 2025; Liu et al. 2025). Gene orthology was predicted using OrthoFinder v2.5.4 (Emms and Kelly 2019). Single-copy orthologs (SCOs) were then aligned using PRANK v.170427 with the '-F' option and trimmed using Gblocks v0.91b with the '-b5=h' option (Castresana 2000; Loytynoja and Goldman 2005). For SCOs: trimmed alignments of Hierarchical Orthologous Groups (HOGs) together with the species tree produced by OrthoFinder were used for the selection analyses with CodeML (PAML version 4.9j), HyPhy RELAX and aBSREL v2.5.41 (Yang 2007; Kosakovsky Pond et al. 2020). CodeML configs had the following parameters: runmode = 0, seqtype = 1, CodonFreq = 2, ndata = 1, clock = 0, model = 1, NSsites = 0, fix omega = 0, omega = .4, cleandata = 0. To test relaxed and positive selection, we used HyPhy RELAX and aBSREL with (i) each branch as foreground and (ii) three branches leading to termites with bifurcated development as foreground and other termite branches as background. To check how many genes under relaxed or intensified selection could be identified by chance, we assigned three random termite branches as foreground 100 times and repeated the RELAX analysis for each iteration. All other remaining HOGs were aligned as above. For each HOG, we reconstructed phylogenies using FastTree v2.1.11 with default options (Price et al. 2010). The *dN/dS* ratios were then reconstructed with CodeML with the same config parameters. Benjamini–Hochberg (BH) corrections were implemented to account for multiple testing.

**MCMCglmm**

As selection patterns can be related to change in ecological and social traits (Rubin 2022), we estimated how *dN/dS* values of SCOs and all remaining HOGs, as well as the parameter *k,* which quantifies the distribution of *dN/dS* values for SCOs, correlated with different measures of sociality and other potentially relevant ecological traits. To account for phylogeny, we applied a phylogenetic generalized linear mixed model based on a Monte Carlo Markov Chain (MCMC) method using the R package “MCMCglmm” v2.34 (Hadfield 2010). The *dN/dS* or *k* parameter values were set as the dependent variable, and the different measures of sociality were set as the fixed effect. Due to the gamma distribution of the dependent variable, we applied the “exponential” link family from the MCMCglmm package to the models. Each model contained only a single fixed effect as follows:

$$ysi = \beta 1 : s + \mu p : s + \mu r : s + \mu o : i + esi \tag{S1}$$

In S1, *ysi* is the selection value for the SCO or all remaining HOGs *i*, for the species *s*. The fixed effect $\beta 1$ represents the social measure or ecological trait tested for each species. The random effect $\mu p$ represents the effect of the phylogeny assuming a Brownian motion model of evolution, the random effect $\mu o$ was applied to account for the variation between orthologous groups, and *esi* represents model residuals. Each species was given a random ID which was specified in an additional random effect $\mu r$. This random effect estimates the non-phylogenetic component of between-species variance. Models for SCO were run for 13 million MCMC generations, sampled every 5000 iterations with a burn-in of 3 million generations. Models for remaining HOGs were run for 1.3 million generations sampled every 5000 iterations, with a burn-in of 300,000 generations. Effective sample size and autocorrelation were verified afterwards to ensure model convergence. Models were fitted with three different uninformative priors of different distributions (parameter expanded, flat, and inverse-Wishart priors) to test the robustness of the models to changes in the degree of belief, *nu*. Inverse-gamma priors were chosen for the residual variances, with a shape and scale equal to 0.001

(Hadfield 2010). The proportion of the between-species variance that can be explained by phylogeny was calculated using the equation *Vp /( Vp + Vr)*, where *Vp* and *Vr* represent the phylogenetic and non-phylogenetic between-species variance, respectively, and are equivalent to phylogenetic heritability or Pagel's lambda (Pagel 1999; Housworth et al. 2004). We report the median of the parameter posterior density, and their 95% credible intervals (CIs) for each explanatory variable tested and the phylogenetic, non-phylogenetic and orthogroup variance. The result of a model is considered significant if CI does not overlap zero. Model results with expanded priors including the non-phylogenetic effect are given in the text, except where otherwise stated.

*OCM* and *SPL* differences (Supplementary text) between FW and TW species were investigated with a simplified version of model S1: *ys* = *OCM* or *SPL* values for species s; $\beta 1{:}s$ represents the social category "Social (FW)" or "Social (TW)" (solitary and subsocial were set to NA due to insufficient sample size. For each species, $\mu p{:}s$ and $\mu r{:}s$ were retained while $\mu o{:}i$ was removed and *es* represented model residuals. Number of iterations were set to 13 million MCMC generations with sampling every 5000 iterations with a burn-in of 3 million generations. Priors, between species variance and model evaluation were as above.

**Gene family size evolution**

We further sought to identify if certain gene families were expanded or contracted at key transitions in termite social complexity, as explored previously in Blattodea (Harrison et al. 2018; He et al. 2021; Mikhailova et al. 2024) and Hymenoptera (Simola et al. 2013; Kapheim et al. 2015). We predicted the expansion and contraction of all orthologous gene families with CAFE5 using the fossil-calibrated termite phylogeny and the HOG counts from OrthoFinder (Mendes et al. 2021). To estimate the best-fitting model for our dataset, we estimated the number of gamma ($\gamma$) rate categories, *K*, which is the number of categories of gene-family-specific rates of evolution, and the number of clusters of branch-specific average rates of gene gain/loss, lambda (λ). Firstly, we estimated the optimal number of gamma rate categories by setting the number of lambda clusters, *L*, to one (i.e. a single average rate of gene gain/loss throughout all branches of the tree) and varying *K* from one to 10. Each model was repeated 50 times to ensure convergence of lambda. We compared AIC values from the models in which lambda converged to a single estimate. Models with up to three *K* categories gave convergent values of lambda among which the lowest AIC value was for the model with *K* = three categories ($AIC_{K=1,L=1}$ = 777538, $AIC_{K=2, L=1}$ = 677512, $AIC_{K=3, L=1}$ = 676803). Secondly, we estimated branch-specific lambda values on the tree by setting the parameter *K* to one and allowing different values of lambda for one branch of the tree. All 48 trees were repeated between 150 and 250 times to ensure convergence of branch-specific lambda values. We grouped branch-specific lambdas in two to six clusters using k-mean clustering with the R packages "cluster" v2.1.4 (Maechler et al. 2013) and "factoextra" v1.0.7 (Kassambara and Mundt 2017). We repeated the models 50 times with varying numbers of gamma rate categories (one to three) and lambda clusters (one to six) to test for convergence of lambda values. Based on the convergence of lambda and AIC values, the model with three gamma rate categories ($\gamma$ = three) and a single cluster of lambda best fitted the data ($AIC_{k=3, L=1}$ = 676803). We used the median value of lambda from the selected model ($\lambda$ = 0.24314831349735) to estimate gene size evolution of all gene families.

**Gene family size differences**

CAFE analysis is restricted to gene families for which copies of the genes are non-zero at the root of the tree. Therefore, of the 33338 HOGs found in the 29 sampled species, only 17438 could be included in the CAFE5 analysis. Of these, 569 gene families showed significant size evolution, although not at the nodes of the transition from linear (FW) to bifurcated (TW) ontogeny. We applied an independent phylogenetic generalized linear mixed model approach (MCMCglmm, see above) using a Gaussian family distribution to examine whether evolution in these gene families was significantly associated with ontogenic patterns of termite sociality. We carried out models with a phylogenetic and non-phylogenetic random effect as well as with a phylogenetic random effect only. We repeated models with two types of priors for the random effects (Inverse-Wishart and expanded priors) and used a Gamma prior for the residuals. Each model was run with every gene family, applying a Benjamini–Hochberg (BH) correction for multiple testing. From the different models, a total of 44 gene families (from N=569 significant HOGs as identified by CAFE5) showed a significant size difference between species with linear (FW) and bifurcated (TW) ontogeny. As we were interested in gene family size evolution in each of the three branches where true workers had convergently evolved, we paid particular attention that the difference was present in all independent transitions to a bifurcated ontogeny. For the 2 true worker branches represented by a single species (*M. darwiniensis* and *D. longilabius*), gene families were only considered when values were closer to the median of termites with bifurcated (versus linear) development. Therefore, from these 44 gene families, only 10 showed a consistent pattern across all three branches. We applied the same approach to the 15900 gene families that could not be analyzed by CAFE5, filtering out gene families with fewer than 13 copies over the tree (corresponding to the number of sampled FW species). Of the 1865 families that passed this threshold, none showed a significant size difference between linear (FW) and bifurcated (TW) species.

**Odorant receptor annotation and phylogeny**

Following the identification of expanded Odorant Receptor (ORs) gene families, we re-annotated ORs in the genomes to further validate our findings. To annotate Odorant Receptors (ORs), we used the software BITACORA v1.3, which performs similarity-based search with BLAST and HMMER to annotate specific gene families not only in the proteome but also in the genome and outputs new gene models (Vizueta et al. 2020). The database for gene annotation included OR sequences from *Diploptera punctata, Blatella germanica, Drosophila melanogaster, Apis mellifera and Apolygus lucorum* (Robertson et al. 2018, Fouks et al. 2023). The Hidden Markov Model (HMM) profile for the database was built with hmmbuild v3.3.2 and mafft –auto v7.397. We ran BITACORA with default settings, except for omitting the length filter and focusing on the presence of domains of interest instead. Curated gene models were filtered for the presence of the 7tm_6 domain by using pfam scan.pl script with default settings (pfam1.6, PFAM database v37.1) and the transmembrane domain by using DeepTMHMM 1.0 (Hallgren et al. 2022). The reconstruction of OR phylogeny was performed according to Johny et al. (2023), as follows. All OR sequences were aligned with mafft-einsi v7.505, then trimmed with trimal v1.4.1 (-automated1 option). The alignment included one GR sequence from *Blattella germanica* as an outgroup. Based on this alignment, a phylogeny was built using IQTREE2 v2.1.3 with the parameter "-m TEST". Visualization and annotation of the phylogenetic tree was done with the R packages: iTOL v7.2.1 and itol.toolkit v1.1.7 (Zhou et al. 2023, Letunic and Bork

2024). Clustering of ORs was performed with Possvm v1.2 (Grau-Bové et al 2021) using the gene tree as input at default settings. To investigate signatures of positive selection, we aligned each OR cluster of interest and reconstructed gene phylogenies as described above. Positive selection was tested using the HyPhy aBSREL v2.5.41 software without specifying the test branches (Kosakovsky Pond et al. 2020).

**Estimated methylation levels of protein-coding genes**

To estimate the methylation levels of protein-coding genes, the metric CpGo/e was calculated by comparing observed to expected CG counts, where expected counts are the product of cytosine and guanine fractions. The distribution of CpGo/e values per species was plotted using the R function "plot" (density(CpGo/e)). Modes were calculated with the function Modes() from the "LaplacesDemon" v16.1.6 package in R. Principal component analysis of CpGo/e values for SCOs was performed with the function prcomp() from the "stats" v4.4.1 package in R. Data visualization was performed with R (base R v4.2.0 and ggplot2 v3.4.4, Wickham 2016).

**Analysis of brain gene expression patterns across 14 species of roaches and termites**

We carried out a comparative evolutionary analysis of brain gene expression patterns across 14 species of termites and cockroaches to examine how brain genetic machinery evolved during the termite MET.

**Insect samples**

Insects used for brain dissection were either collected in the field or reared in breeding rooms. *Blattella germanica* and *Blatta orientalis* were maintained at the Federal Institute of Materials Research and Testing (BAM), Berlin, Germany at 28 °C and subjected to a 12 h:12 h dark/light cycle. Their diet consisted of a mixture containing 77.0% dog biscuit powder, 19.2% oat flakes and 3.8% brewer's yeast and supplied with water. Oothecae were individualised, and family-groups of cockroaches were bred and instar development was followed until adult emergence. In each family group, we randomly sampled one female and one male of the 4th nymphal instar for *B. germanica*, one female and one male of the 6th nymphal instar for B. orientalis, and one male and one female adult for both species. Five family groups were pooled in one replicate, and no family group was used twice in a different replicate.

*Cryptocercus meridianus* was collected in Yunnan, China in May 2016 (He et al. 2021) and stored in RNA-later until dissection. *Cryptocercus punctulatus* was reared at the National Natural History Museum (MNHN), Paris, France. *Mastotermes darwiniensis* and *Prorhinotermes simplex* were maintained at the BAM at 29°C, 80% rH, *Reticulitermes flavipes*, *Neotermes castaneus*, *Kalotermes flavicollis* and one colony of *Coptotermes gestroi* were kept at 28°C 70% rH. All termites were kept in total darkness. Two colonies of *C. gestroi* were reared and dissected at the Sao Paulo State University (UNESP), Rio Claro, Brazil. *Anoplotermes pacificus* colonies were collected in Sao Paulo, Brazil and dissected at the Federal University of ABC (UFABC), Sao Paulo, Brazil. *Zootermopsis nevadensis* and *Hodotermopsis sjostedti* were reared and dissected at OIST, Okinawa, Japan. One colony of *H. sjostedti* was reared at the Universite Sorbonne Paris Nord, Paris, France. *Macrotermes natalensis* was collected in Pretoria, South Africa, by David Sillam-Dussès. Alates were not available for the same colonies as workers and soldiers in this species. Caste and sex identification were carried out by examination of external structure. The shape of the last tergites was examined for sex determination of non-Termitidae. For the two species of Termitidae where morphological sex determination is not possible, sex determination was done based on body size polymorphism for *M. natalensis* (Noirot 1955). Worker sex determination of *A. pacificus* was not carried out and both sexes were pooled. When possible, 5 colonies were pooled per sample. For *C. meridianus, C. punctulatus, M. darwiniensis, Z. nevadensis, H. sjostedti, C. gestroi, and M. natalensis*, only one colony per sample was used. All samples were kept alive before dissection except for *C. meridian*us that was conserved in RNA-later at -80°C. For *C. punctulatus, M. natalensis* and half the samples of *B. germanica* and *B. orientalis*, samples were snap frozen and conserved at -80°C until dissection. When not mentioned otherwise, dissections and RNA extractions were carried out at BAM and by the same person.

**Brain dissection**

We carried out brain dissection on termites and cockroaches. When possible, samples were alive prior to dissection, if not, they were kept at -80°C until dissection and thawed slowly. Insects were anaesthetized when necessary and swiftly beheaded. The head was directly placed in ice-cold Phosphate Buffer Solution (PBS) and dissected under a stereoscope (Olympus). The cuticle of the head capsule was punctured above the brain region and removed to access the brain. Adipose tissue surrounding the brain area was removed and the brain was gently pulled out of the cephalic capsule. Brains were snap-frozen in liquid nitrogen and stored at -80°C until RNA extraction. All dissections were performed below 4°C and within 10 minutes after beheading. A total of 1492 brain dissections were performed. To reduce colony- and individual-based variation and ensure sufficient RNA quantity, each RNA sample contained a pool of up to 5 brains, except for *A. pacificus* workers where 10 brains were required. A total of 285 RNA samples were generated.

**RNA extraction and sequencing**

Total RNA was extracted using the RNeasy Plus Mini Kit (Qiagen, Germany) including a gDNA eliminator column as per the manufacturer's protocol, which provided higher yields with brain tissue. RNA was quantified on a Qubit (Invitrogen) and RNA integrity was assessed on an Agilent 2100 Bioanalyzer (median RIN for all samples 8.5). RNA was then stored at -80°C until library preparation. Messenger RNA (mRNA) enrichment was performed using the NEBNext® Poly(A) mRNA Magnetic Isolation Module, cDNA synthesis and further library steps were carried out with the NEBNext® Ultra™ II Directional RNA Library Prep Kit for Illumina®, and paired-end 2x100 bp sequencing was performed at a depth of 50 M reads per sample on the NovaSeq 6000 with a S4 XP v1.5 Flowcell at the DRESDEN-concept Genome Center (DcGC).

**Read pre-processing**

We generated a total of 285 RNA-seq datasets from 10 species of termites and 4 species of cockroaches covering representative termite caste diversity and the main life stages of cockroaches. For each stage or caste, we generated transcriptomic data for both males and females, with at least 3 replicates for each when possible. All replicates were first trimmed using TRIMMOMATIC v0.39 with ILLUMINACLIP:TruSeq3-PE-2.fa:2:30:10 and default parameters (Bolger et al. 2014). We used HISAT2 v2.1.1 (Kim et al. 2019) to map the reads on their respective genomes, indexed beforehand with hisat2-build. Aside from *C. meridianus* whose RNA quality was low (average RIN 4.95), and one sample of male nymphoid of *R. flavipes*, the overall alignment rate was above 70% for all species. The resulting SAM files were successively sorted by name using SAMtools v1.17 (Li et al. 2009). We computed read counts using htseq-count v2.0.9 (Anders et al. 2015; Putri et al. 2022) in combination with the name-sorted SAM files and the gff3 annotation of each respective genome. We used default parameters but for the stranded parameter set to 'no', the feature type set to 'gene', and the id attribute set to 'ID' to extract the read count per gene. For each species, we generated a read count table with the gene IDs in rows and sample in columns as well as a condition file indicating the species name, the caste or stage name, the sex and the replicate ID for each sample. For each of the 14 species of cockroaches and termites, read counts were normalized using the median of ratios from DESeq2 (Love et al. 2014). SAM files were converted into BAM files and stored with the raw reads and the gene counts.

**Gene co-expression network construction**

We generated differential networks for each species through a full comparison of caste-specific networks to identify genes involved in the regulation of caste determination. Within each species, normalized read counts were grouped per caste to calculate caste-specific gene co-expression networks. Male and female samples were always balanced within stages or castes (except for sex-specific castes) and so were merged to increase the number of samples per gene co-expression network. We calculated the signed values of the weighted topological overlap (WTO) for each caste of each species using the R package "wTO" v2.0.2 (Gysi et al. 2018). WTOs for each link of a network were computed including all genes present in the species transcriptome, without bootstrapping. A total of 34 caste-specific networks were produced for the 10 termite species and 8 stage-specific networks for the 4 cockroach species. Because all genes were included in the analysis, all networks within a species had the same number of nodes and links. We estimated the caste specificity of the genes included in each species network using the R package "CoDiNA" v 1.1.2 (Morselli Gysi et al., 2020). We provided CoDiNA with all caste-specific networks for a given species and ran network comparisons to generate differential species networks. In this network, CoDiNA infers a caste-specificity for each link. If a link is attributed to a caste, it means that this link is only present in this particular caste. Next, the genes were attributed to a caste based on the caste specificity of the links. For example, if a gene had a majority of links attributed to a worker caste, the gene was assigned to a worker-specific network. We only considered caste-specific genes if these were attributed to a single caste (i.e. not specific to more than one caste). We extracted the list of genes that were worker-, soldier-, secondary reproductive- (i.e. including neotenic, ergatoid, and nymphoid caste), nymph-, juvenile-, adult-, primary reproductive- (i.e. alate and king and queen) specific. The lists of genes from these categories are hereafter called the list of caste-specific genes (full comparison). Note that their specificity comes from their connection in the network, and therefore are more likely to be involved in caste-regulation. All network comparisons with CoDiNA were performed using default parameters and node clustering was performed with median quantiles for the internal and external cutoffs which offer more permissive values.

A second set of differential networks were generated via pairwise comparison of worker and reproductive networks for all 4 cockroach species and 9 species of termites. *Zootermopsis nevadensis* was excluded from the pairwise comparisons due to the absence of a reproductive caste for this species. Worker and reproductive networks were computed as described above and compared in a pairwise manner, again using CoDiNA. As before, we only considered genes to be caste-specific if these were attributed to one of the two compared castes (i.e. worker or reproductive) and extracted corresponding gene lists. The genes from these categories are hereafter referred to as pairwise caste-specific genes. Again, all network comparisons with CoDiNA were performed using default parameters and node clustering was performed with median quantiles for the internal and external cutoffs.

**Gene co-expression network visualization**

Reduced gene co-expression networks containing ORs and DEGs were visualized with Cytoscape v3.10.3. Edge widths were scaled by absolute correlation coefficient, only including those with an absolute correlation coefficient greater than 0.4. Nodes with 5 edges or more were included, and were colored and shaped according to the DEG or OR category. The network layout was generated with the prefuse force directed method, weighted by absolute correlation coefficient.

**Differential gene expression**

Normalized read counts were used to analyze DEGs between castes to search for evidence of a genetic toolkit associated with different levels of termite social complexity. We performed differential gene expression in pairwise comparison between castes of the same species using DESeq2 (Love et al. 2014). Model design only included caste as a fixed effect and male and female samples were merged. Read counts were normalized using the median ratios. Non-expressed genes were removed from the analysis. A gene was considered differentially expressed if the Benjamini-Hochberg (BH) adjusted p-value was below 0.05. Due to variabilities in differential expression and the high number of species comparisons, the minimum Log fold change for DEGs was set to zero. For each pairwise comparison, DEGs were classified as caste-biased in one of the two castes being compared. We searched for caste-biased genes by searching for overlaps of orthologue IDs across species using the lists of caste-biased genes from pairwise comparisons. For example, after identifying worker-biased genes from comparisons with reproductives in all 9 species with false or true workers (*Z. nevadensis* was not included in such pairwise comparison due to the lack of reproductive samples), we compared those lists to identify the worker-biased orthologue ID overlaps.

**Principal Component Analysis (PCA)**

Normalized read counts were used to perform PCAs and verify if samples clustered by caste or sex. For each species, we performed a PCA for every species with the R package FactoMineR (Le et al 2008) and factoextra (Kassambara and Mundt 2017). For each PCA, we explored the partitioning of the cluster based on caste (or stage), sex, and replicates over the first 5 principal components. Replicate clusters overlapped in all species except species where colonies originated from two different locations (*H. sjostedti* and *C. gestroi*) or were conserved in two different ways before dissection (*B. orientalis, B. germanica*). However, in all cases, caste clustering was not affected by replicate identity. We further sub-selected from the normalized read counts all SCOs that were biased in pairwise comparison between workers (or juveniles) and reproductives for 13 species (again excluding *Z. nevadensis*). For each species, we averaged the read counts per caste and then averaged and scaled each average gene read count across caste. As such we were able to correct for differences between species and thereby represent all species in the same PCA. We produced such species-normalized PCAs with all SCOs that were identified to be biased in pairwise comparisons between workers, soldiers and reproductives for all 9 termite species (*Z. nevadensis* was excluded) for termite species with bothlinear (FW) (N=4) and bifurcated (TW) (N=5) ontogeny.

**Co-expression interactions between ORs and DEGs**

To investigate the relationship between ORs and DEGs, we generated gene co-expression networks comprised of all ORs and significant worker- and reproductive-biased DEGs, resulting in a worker and a reproductive-network for each termite species (N=9). Caste-specificity of ORs was determined by pairwise comparison of these worker and reproductive co-expression networks. We considered only the DEGs that were shared by all 5 TW and all 4 FW species, including both worker and reproductive biased genes. These DEGs are referred to hereafter as shared DEGs. Using the Netvis function of the wTO package, we calculated the number of links between worker- and reproductive-specific ORs and worker- and reproductive-shared DEGs in both worker and reproductive networks (excluding all among-OR and among-DEG links from the count as well as all links below a wTO threshold of 0.33). The number of connections was then divided by the number of ORs and then by number of DEGs, resulting in a proportion of links between ORs and shared DEGs for each species. We performed this analysis for all ORs, gene cluster 1-ORs and gene cluster 2-ORs (see main text). This resulted in proportions of connections between worker- or reproductive-specific ORs with worker- or reproductive-biased shared DEGs in the worker- or reproductive- network of every species.

**Selection and TE content of DEGs**

We examined the following properties of caste-biased genes: (i) *dN/dS* ratios for multi-copy orthologs identified as described above, (ii) *k* values for SCOs as identified and as described above, and (iii) number of transposable elements in the 10kb upstream region of a gene (TEs identified in Liu et al. 2025, counted using the bedtools (v2.30.0) intersect). These properties were compared between worker- and reproductive-upregulated genes in termites with linear and bifurcated development as well as juvenile- and adult-upregulated genes in solitary and subsocial cockroaches, the latter two of which were combined into a single category for the TE analysis, due to availability of TE annotations from only one solitary (*B. orientalis*) and one subsocial (*C. meridianus*) species (Liu et al. 2025). We used generalized linear mixed models implemented in the package lme4 in R to evaluate the relationship between the gene properties as follows:

$$y = Group * ExprBias + 1|Species \tag{S2}$$

In S2, $y$ is one of the gene properties, for example the *dN/dS* ratio, $Group$ is the social category (solitary cockroaches, subsocial wood roaches, linear (FW) and bifurcated (TW) termites. $ExprBias$ is a caste or stage category in termites or cockroaches, respectively. Species ID was used as a random variable. A comparison of different groups in terms of development and expression bias was performed using the "emmeans" package in R (v1.10.0).

**GO term analysis and orthologue group comparisons**

We identified GO-term enrichment of differentially expressed genes as well as the caste-specific genes identified via the network analysis. For each species, we retrieved domain annotations of the longest isoforms with PfamScan. We mapped the pfam IDs to GO-terms based on the Mitchell et al. (2015) pfam to GO documentation and generated TopGO-compatible universe files for each species using custom-made scripts. GO universe and gene lists of interest were used in TopGO v.2.50.0 (Alexa and Rahnenfuhrer 2022). TopGO objects with Biological Process and Molecular Function ontogenies were generated using "annFUN.gene2GO" annotation function and a custom-made "GOmap" script. We performed significance tests on GO-terms of the GO-object using the "weight01" algorithm and "fisher" statistical test. We retrieved all GO terms that were significantly overrepresented (at alpha = 0.05) than expected by chance. In the network analysis, GO-term analyses on common genes were computed to ensure that general housekeeping functions were represented. Finally, we extracted the gene names contributing to the enriched GO-terms.

**Identification of genetic toolkits**

We used the R package "SuperExactTest" v1.1.0 (Wang et al 2015) to test the significance of overlapping sets of orthologs between castes and across species. After carrying out pairwise comparisons of DEGs between worker (or juvenile), soldiers and reproductives, we extracted the HOG ID of the genes. For each pairwise comparison we identified the HOGs that were overlapping among species. We reported HOGs that were i) worker- and reproductive-biased in 9/9, 7/9, 5/9 termite species comparisons, ii) soldier and reproductive biased in 8/8, 6/8 and 4/8 termite species comparisons, and iii) worker- and soldier-biased in 9/9, 7/9, 5/9 termite species comparisons. We additionally reported HOGs that were juvenile and adult biased in subsocial cockroaches (2/2) and solitary cockroach (2/2). To further compare TW and FW species, we searched for overlapping HOGs among all 4 linear (FW) species (5 in soldier versus worker comparisons) and all 5 bifurcated species (in addition to 4 species including *M. darwiniensis*). To calculate whether an overlapping HOG across multiple species was not due to random chance, we performed significance tests by sub selecting only HOGs that had at least one orthologous gene across the species comparisons, with the disadvantage that some HOGs could not be tested for significance. For example, in comparisons of all 9 termite species, a candidate caste-biased HOG could be shared by 8 species, but if it was only present in the genomes of those 8 species (and not all 9 species), no p-value on the significance of the overlap could be determined. The same method was used to identify overlapping HOGs in caste-specific gene networks as well as pairwise comparisons of worker versus reproductive gene co-expression networks.

**Gene functional annotation**

Reciprocal best blast hits were obtained for all annotated proteins of each analyzed species against the proteomes of *Drosophila melanogaster*, *Apis mellifera*, and *Tribolium castaneum* to provide insights into the potential functions of the identified proteins This was carried out with blastp v.2.12.0+. GO term annotation was produced as described above (see *GO term analysis and orthologue group comparisons*). We extracted the ID of the OG that was differentially expressed between the worker and reproductive caste (or was caste-specifically expressed) and shared between 80% and 100% of the species studied. To identify gene function by homology, we first applied reciprocal blast searches against *Drosophila melanogaster*. If the reciprocal blast did not retrieve any match we used the best blast hit using *D. melanogaster*. If this did not give a majority consensus hit, we checked the reciprocal blast and best blast hit for *Apis mellifera* and *Tribolium castaneum*. In the case of disagreement or if the latter searches did not provide a consensus hit, we classified the orthogroup as “unclear”.  Using this list of genes, we identified JH-related genes, 20E-related genes and Insulin-related genes based on Hanada et al 2025. We categorized the remaining genes based on their putative function: Peptidase, neuronal function (e.g. neurotransmitter, neuron development), other hormone (non-JH), immunity/ROS, gene regulation (including histone and transcription factors), Cytochrome P450, Vitellogenin, and Other (including genes not assigned to the above categories as well as uncharacterized or unclear genes).

In this manner, genes pertaining to HOG001851 (with TW-specific expansions) were identified as being related to CCHamide receptors, as the majority of best blast hits in *D. melanogaster* belonged to CCHa1-R and CCHa2-R. All sequences from this HOG were then aligned and a tree was generated using the same methods applied for ORs. For this gene tree, CCHamide1 receptors were used as a root, that were identified via reciprocal best blast hit with the CCHa1-R gene (FBpp0086052) in *D. melanogaster*.

**Figure formatting**

Multi-panel figures appearing in the main text were assembled and formatted in Inkscape v.1.3.2.

### Supplementary Text

#### Characterization of termite ontogeny

Termite ontogeny boasts a wide diversity of hemimetabolous developmental pathways and has been widely studied to understand the development of diverse termite caste patterns (Noirot 1955; Kaiser 1956; Buchli 1958; Pasteels 1965; Renoux 1976; Watson et al. 1977; Luykx 1993; Roisin 2000; Bourguignon et al. 2009; Moura et al. 2011; Neoh and Lee 2011; Rasib and Saeed Akhtar 2012; Chouvenc and Su 2014; Revely et al. 2021). We retrieved information on the developmental pathways of termites and cockroaches from the literature. Developmental pathways are traditionally represented as a directed network where the nodes represent developmental instars and the links represent the development from one instar to another. However, there is no standardized representation of termite ontogenies and some network representations include a mixture of partial as well as confirmed ontogenies, whereas others represent only the ontogeny of the worker line and some attribute different link types depending on the maturity of the colony (Noirot 1955; Buchli 1958; Chouvenc and Su 2014). For example, incipient colonies often require a reduced number of instars to reach a functional worker or soldier stage compared to mature colonies, as is the case in *Kalotermes flavicollis* (Grassé and Noirot 1958), *Prorhinotermes simplex* (Hanus et al. 2006), *Reticulitermes flavipes* (Buchli 1958) or *Coptotermes formosanus* (Chouvenc and Su 2014). Certain developmental paths only occur under experimental settings or constrained conditions. Caste development responds to social and environmental variation. Alate, neotenic and soldier emergence is controlled by caste ratio and/or the presence of another caste (Myles 1999; Mao and Henderson 2007; Miura and Maekawa 2020). Furthermore, the developmental pathway can be sex-specific in termites and cockroaches (Noirot 1955; Nalepa 1984; Short and Edwards 1991). To study ontogenies comparatively, we included only those networks confirmed for mature termite colonies under natural social and ecological variations. Male and female ontogenies were reconstructed separately. Among the 29 species of Blattodea that were studied here, sixteen species are well documented, others have only partial or no information regarding their development. Where possible, we used the ontogeny of the given species at a mature colony stage (i.e. 5 years after the establishment of the mating pair, Chouvenc and Su, 2014). Sister species (N=10) or close relatives from the same family (N=3) were used when no information was found on the species or the genus, respectively. For each species, we described the ontogeny for males and females to account for sex-specificities. Secondary reproductives were classified into: i) apterous neotenics emerging from pseudergates (false workers), ii) brachypterous neotenics originating from pseudergates, iii) ergatoids i.e. apterous neotenics emerging from true workers, and iv) nymphoids i.e. brachypterous neotenics emerging from the nymphs of termites with a bifurcated ontogeny. Adultoids, alates that do not leave the nest, but which participate in the reproduction of the colony were not considered. The occurrence of secondary reproductives and soldier polymorphism in certain species is documented without the description of their ontogenetic history (Myles et al 1999). Therefore, when the most commonly derived instar from secondary reproductives or pre-soldiers was unclear, the connections were assigned based on the nearest known species.

Shortest characteristic path length (*SPL*), terminal castes and Ontogenic Complexity Metric (*OCM*)

We estimated ontogenic complexity for each species as a quantitative proxy of termite sociality, based on the rationale that each caste has a differentiated function in the colony, and therefore higher caste diversity should represent a higher degree of task specialization and therefore division of labor (Revely et al. 2024). We also expected that a species with a shorter instar number would reach a terminal stage and should begin specific tasks sooner, as shown by two independent mechanisms; the development of soldiers from earlier instars as mentioned above, or by the shortening of instar duration as in Termitidae (Noirot 1955; Chouvenc and Su 2014). In other words, long and/or linear developmental pathways leading to fewer terminal stages responsible for all life history tasks (foraging, reproduction, defense, grooming, nurturing etc.) exhibit relatively lower division of labor, and therefore social complexity, than species with fewer instar stages and/or bifurcated ontogenies leading to many terminal stages. To test this prediction, we measured the shortest characteristic path length (*SPL*) (Boccaletti et al. 2006) in our standardized ontogenetic representations and correlated this with other measures of social complexity such as the number of castes, colony size and nesting type. We calculated the *SPL* value of the developmental pathway (Boccaletti et al. 2006) as a way of quantifying ontogenetic bifurcations and branch lengths for each species. The *SPL* value measures the average shortest path from one instar to another in a network and was calculated by measuring the average number of molts between all instars based on standardized ontogenetic schemes. As the networks are bidirectional, this measure also accounts for a regressive molt. The terminal stages considered were: true workers, soldiers, ergatoids, apterous neotenics, nymphoids, brachypterous neotenics, primary reproductives, minor (small) and major (large) workers as well as minor (small) and major (large) soldiers. Intercastes and castes emerging from parasitism or laboratory manipulation were not considered. We defined a terminal state as one that is not able to moult into a new stage. In the case of pseudergates/false workers, individuals moult into secondary reproductives or alates in most cases (Nagin 1972; Lenz et al. 1982; Luykx 1993; Roisin 2000; Roisin and Korb 2011), are functionally more aligned with larvae and nymphs (Korb and Hartfelder 2008; Roisin and Korb 2011), and therefore do not constitute a terminal stage. In the case of true workers, the proportion of individuals molting into soldiers or ergatoids (where present) in an entire colony is small: in *Mastotermes darwiniensis*: up to 10% of the colony is composed of soldiers, while 0.003% consist of worker-derived ergatoids. Corresponding values for the following genera are: *Coptotermes* spp.: 10% (soldiers) and 0.004% (ergatoids), *Reticulitermes* spp.: 2% (soldiers) and 0.006% (secondary reproductives), *Microcerotermes* spp.: 2% (soldiers) and 0.2% (ergatoids), *Amitermes* spp.: 5% (soldiers) and 0.2% (ergatoids) and *Nasutitermes* spp.: 26% (soldiers) and 0.7% (ergatoids) (Haverty 1977; Myles, 1999). The overwhelming majority of true workers therefore never molt into another (reproductive) caste, thereby constituting a functionally terminal developmental stage, corresponding to the functionally sterile status of these individuals (Revely et al. 2024).

**Fig. S1.** Ancestral reconstruction of workers over the termite phylogeny (Hellemans et al. 2024). Black: FW. Red: TW. Likelihoods are represented as pie graphs at each node.

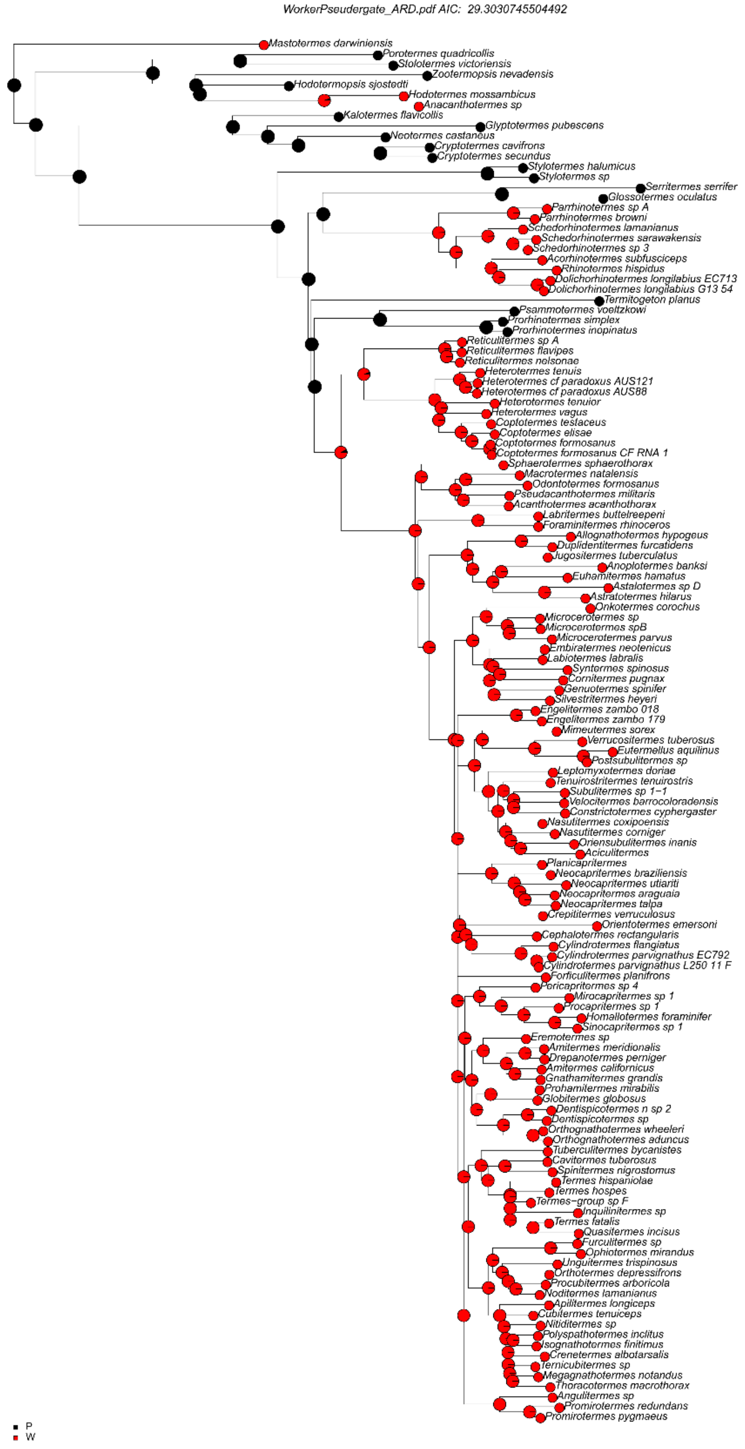

**Fig. S2.** CAFE5 analysis of gene family expansions and contractions over a time-calibrated termite phylogeny. The 3 identified significant expansions are indicated at the corresponding nodes. Notation: Expansion/Contraction/Change per MY).

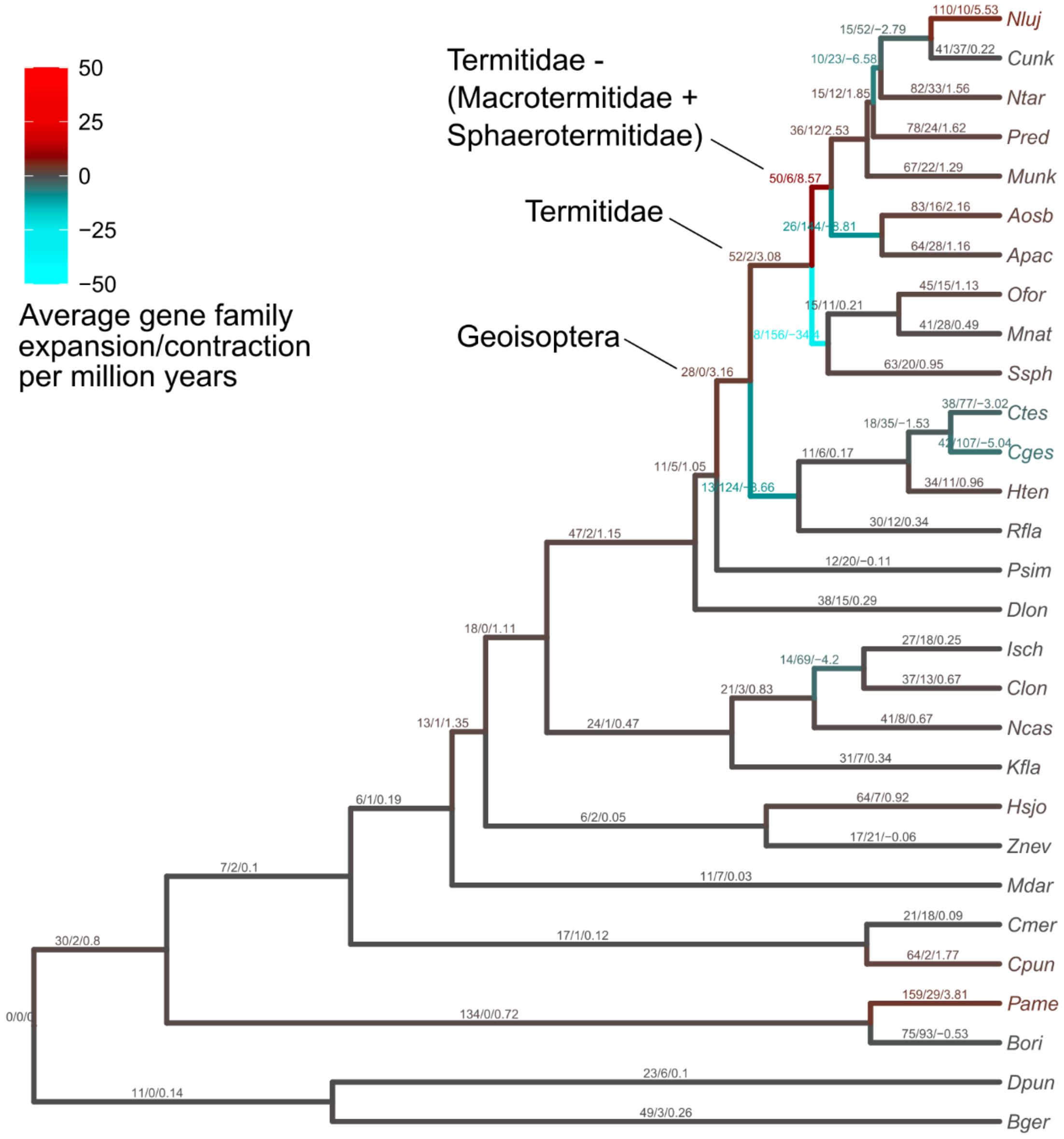

**Fig. S3.** Significant gene family expansions at independent TW transitions under different MCMCglmm models (see Table S2 for details). Gene counts were tested against ontogeny (linear vs. bifurcated) or social category (solitary, subsocial, social (FW), social (TW)). Families were only considered if *M. darwiniensis* (Mdar) and *D. longilabius* (Dlon) clustered more closely with bifurcated (TW) than linear (FW) species. Five families (HOG0000380, HOG00001513, HOG0001851, HOG0000380, HOG0002943) showed significant expansion (EXP) and five families (HOG0000937, HOG0004986, HOG0002046, HOG0002649, HOG0000758) showed significant contractions (CON) in bifurcated (TW) species. Of these, only 2 expansions (HOG0000380, HOG0001851) were supported across all models.

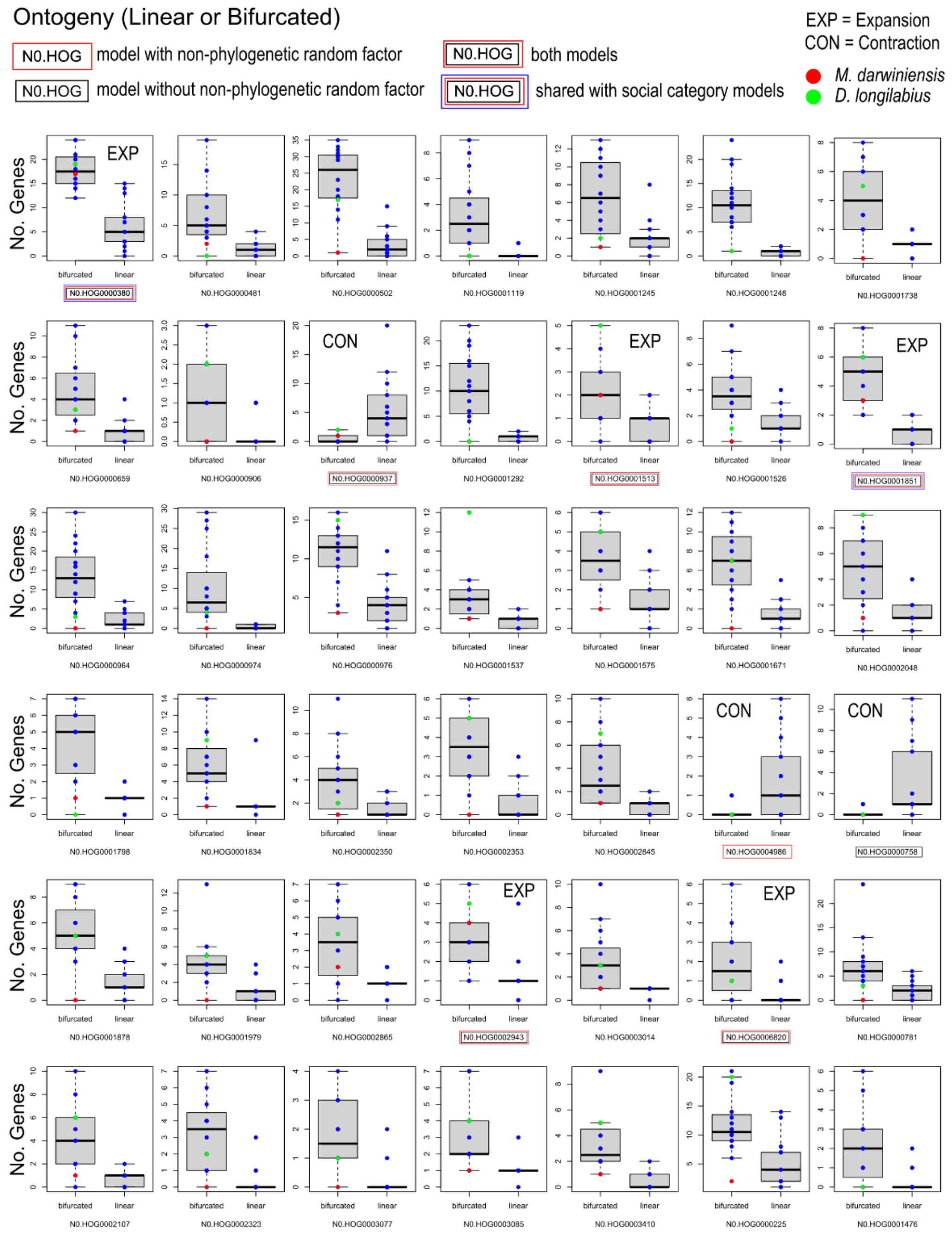

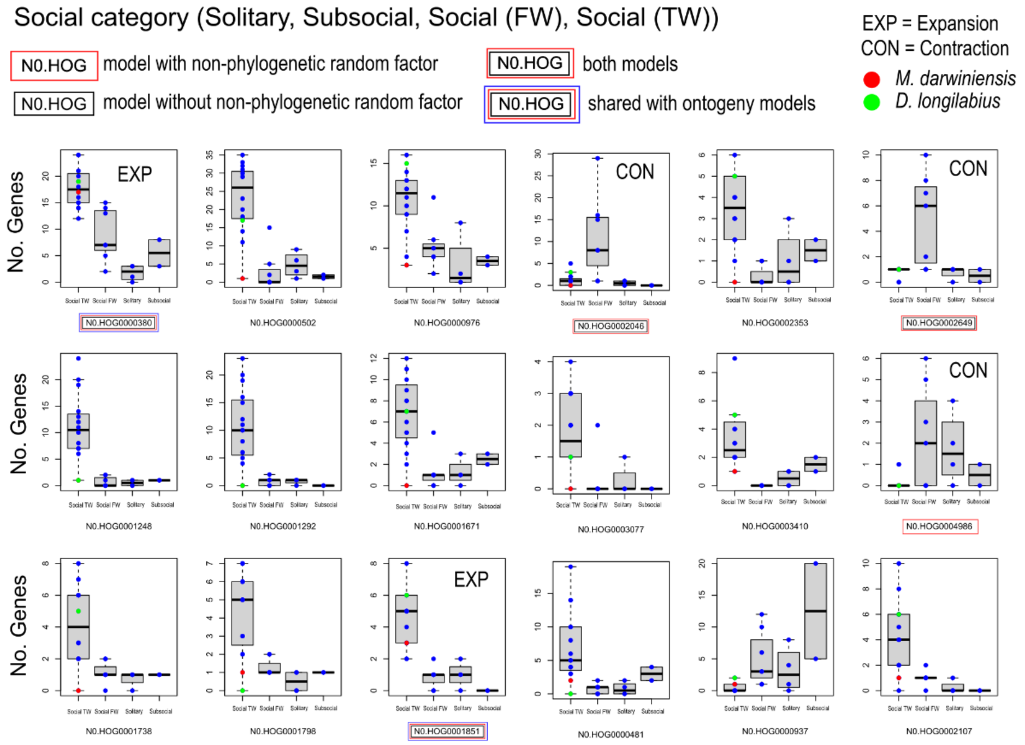
Social category (Solitary, Subsocial, Social (FW), Social (TW))
N0.HOG model with non-phylogenetic random factor
N0.HOG model without non-phylogenetic random factor
N0.HOG both models
N0.HOG shared with ontogeny models
EXP = Expansion
CON = Contraction
M. darwiniensis
D. longilabius
No. Genes
EXP
CON
N0.HOG0000380
N0.HOG0000502
N0.HOG0000976
N0.HOG0002046
N0.HOG0002353
N0.HOG0002649
N0.HOG0001248
N0.HOG0001292
N0.HOG0001671
N0.HOG0003077
N0.HOG0003410
N0.HOG0004986
N0.HOG0001738
N0.HOG0001798
N0.HOG0001851
N0.HOG0000481
N0.HOG0000937
N0.HOG0002107
Social TW
Social FW
Solitary
Subsocial

**Fig. S4.** Gene tree of independent TW-specific expansions to HOG001851, a G-protein coupled receptor (GPCR) sharing sequence similarity with CCHamide brain-gut neuropeptide receptors. The tree is rooted with CCHamide-1-R sequences, which were found as conserved single copies in cockroach and termite genomes, pointing to CCHamide-2 receptors as potential candidates for these genes, which also occur as multiple copies in other insect lineages. Tip branches and labels are coded by social category as in Fig. 3.

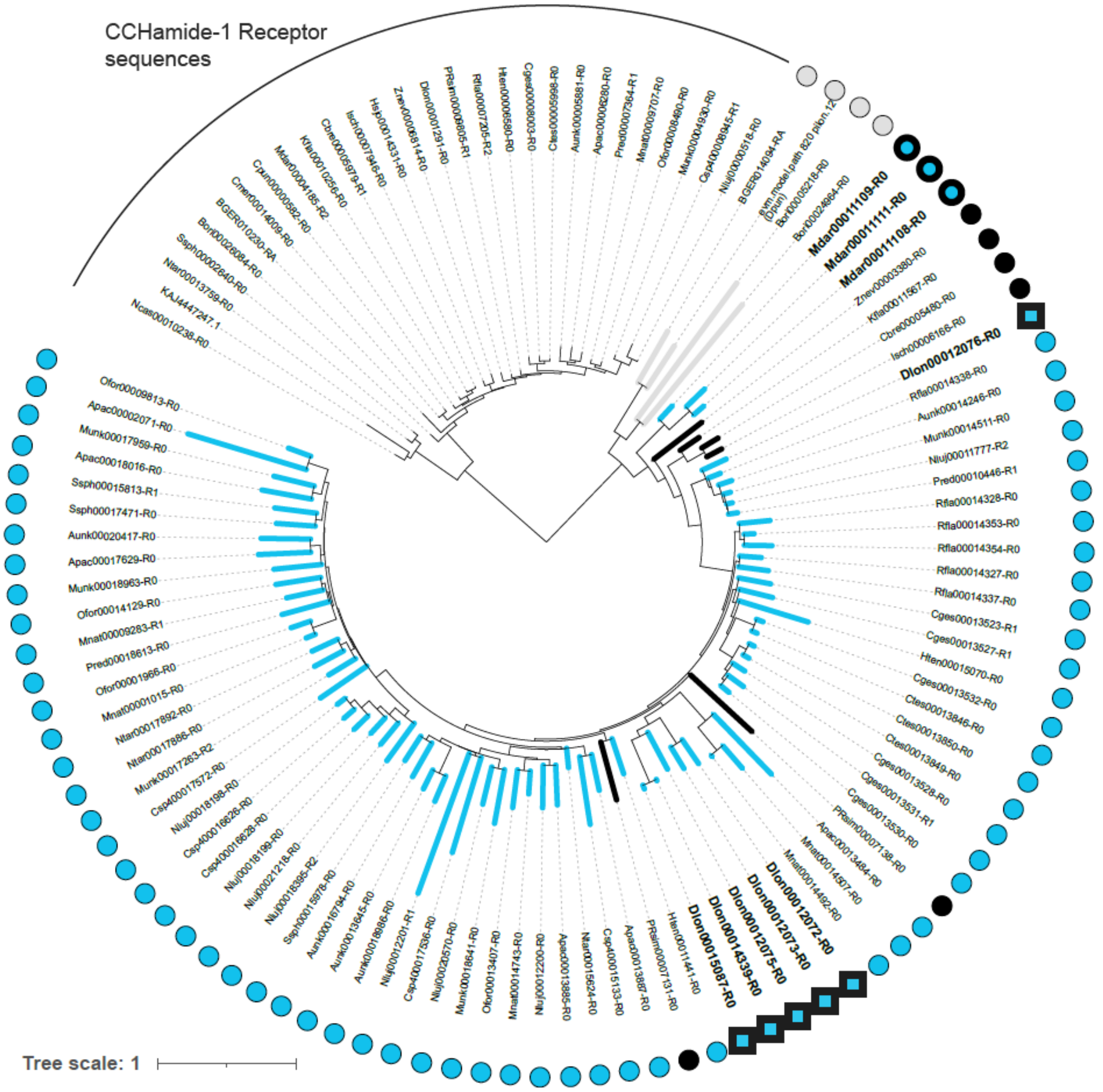

**Fig. S5.** Caste-specific gene co-expression networks highlighting ORs and their connectivity to shared caste-biased DEGs. Left: reproductive networks. Right: worker networks. Nodes represent individual genes. Green: ORs (triangle = gene cluster 1, diamond = gene cluster 2, square = other ORs). Dark blue: reproductive-biased (TW species) DEG. Light blue: TW-biased DEG. Black: reproductive-biased (FW species) DEG. Grey: FW-biased DEG. Edges indicate significant co-expression, with thickness and color representing strength and direction of co-expression, respectively.  Red: negative. Blue: positive. To improve visualization for *Apac* (worker), an edge weight threshold of 0.6 and node degree of 32 was applied. *Mdar = M. darwiniensis*, *Hsjo = H. sjostedti*, *Kfla = K. flavicollis*, *Ncas = N. castaneum*, *Psim = P. simplex*, *Rfla = R. flavipes*, *Cges = C. gestroi*, *Mnat = M. natalensis*, *Apac = A. pacificus*.

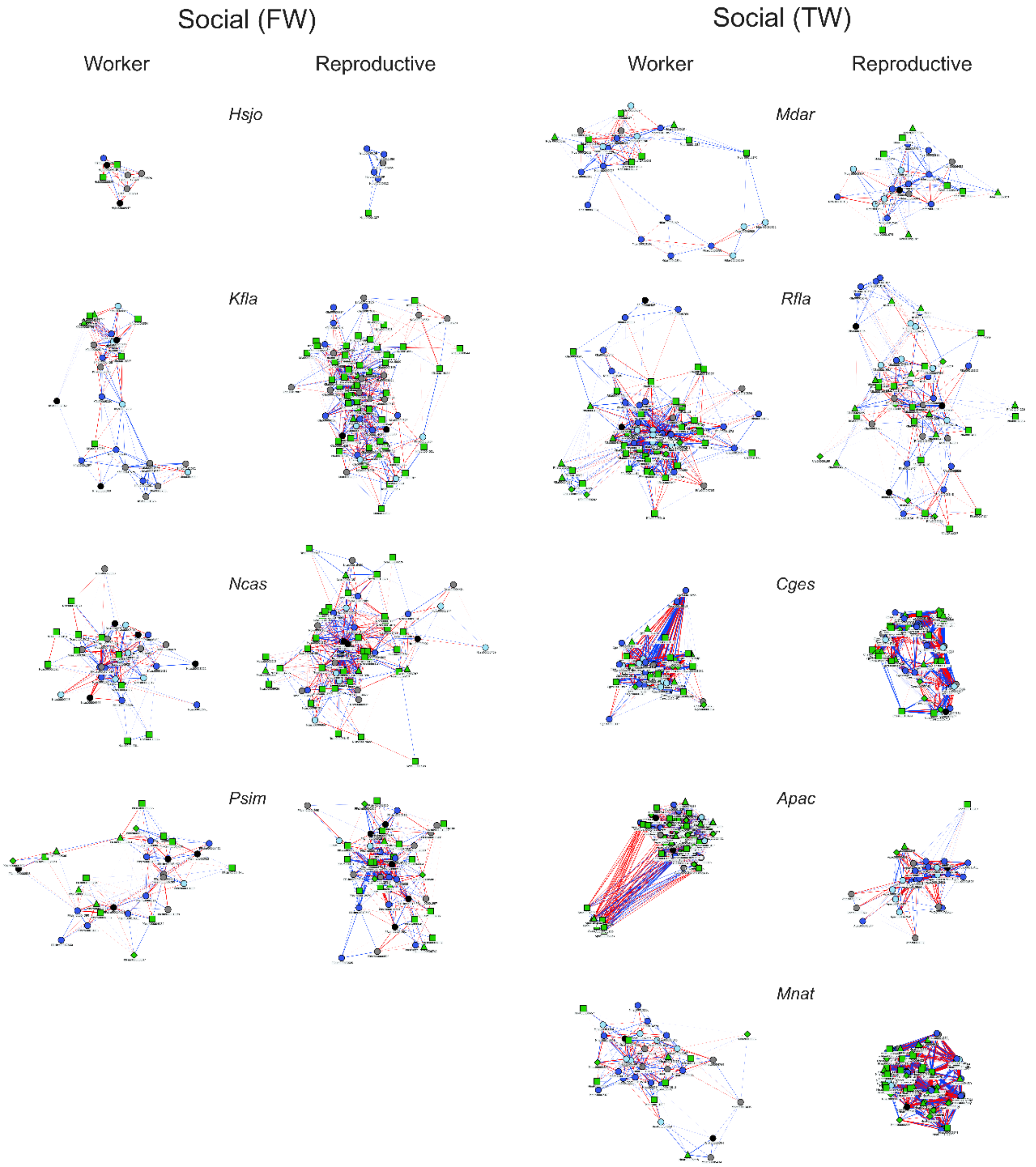

**Fig. S6.** PCA of species-normalized gene expression patterns by sex (left) and species (right). *Mdar* = *M. darwiniensis*, *Hsjo* = *H. sjostedti*, *Kfla* = *K. flavicollis*, *Ncas* = *N. castaneum*, *Psim* = *P. simplex*, *Rfla* = *R. flavipes*, *Cges* = *C. gestroi*, *Mnat* = *M. natalensis*, *Apac* = *A. pacificus*. *Mnat* is not included due to some castes being determined by sex. *Apac* is not included in the lower panels to enable better visualization of other species' data points.

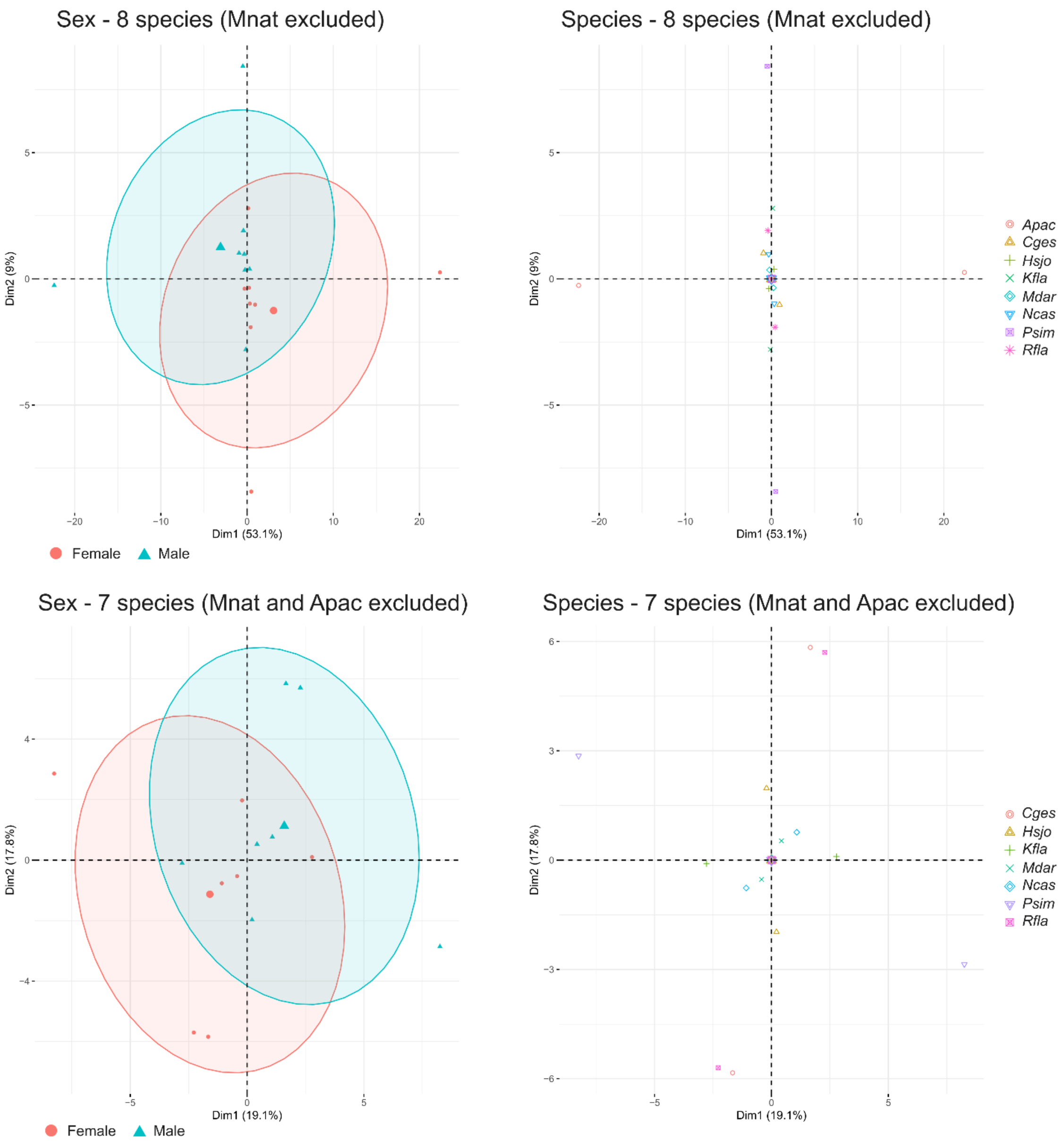

**Fig. S7.** Per species (a) TE content and (b) *dN/dS* ratios of DEGs by simplified social category (all cockroaches (solitary/subsocial), linear (FW) and bifurcated (TW) termites) divided by caste-bias (cockroach: Ad – adult, Ju – juvenile, None – non-biased; termites: Repr – reproductive, Wo – worker, None – non-biased). For the TE analysis, only species were included that had been TE-annotated by Liu et al. (2025).

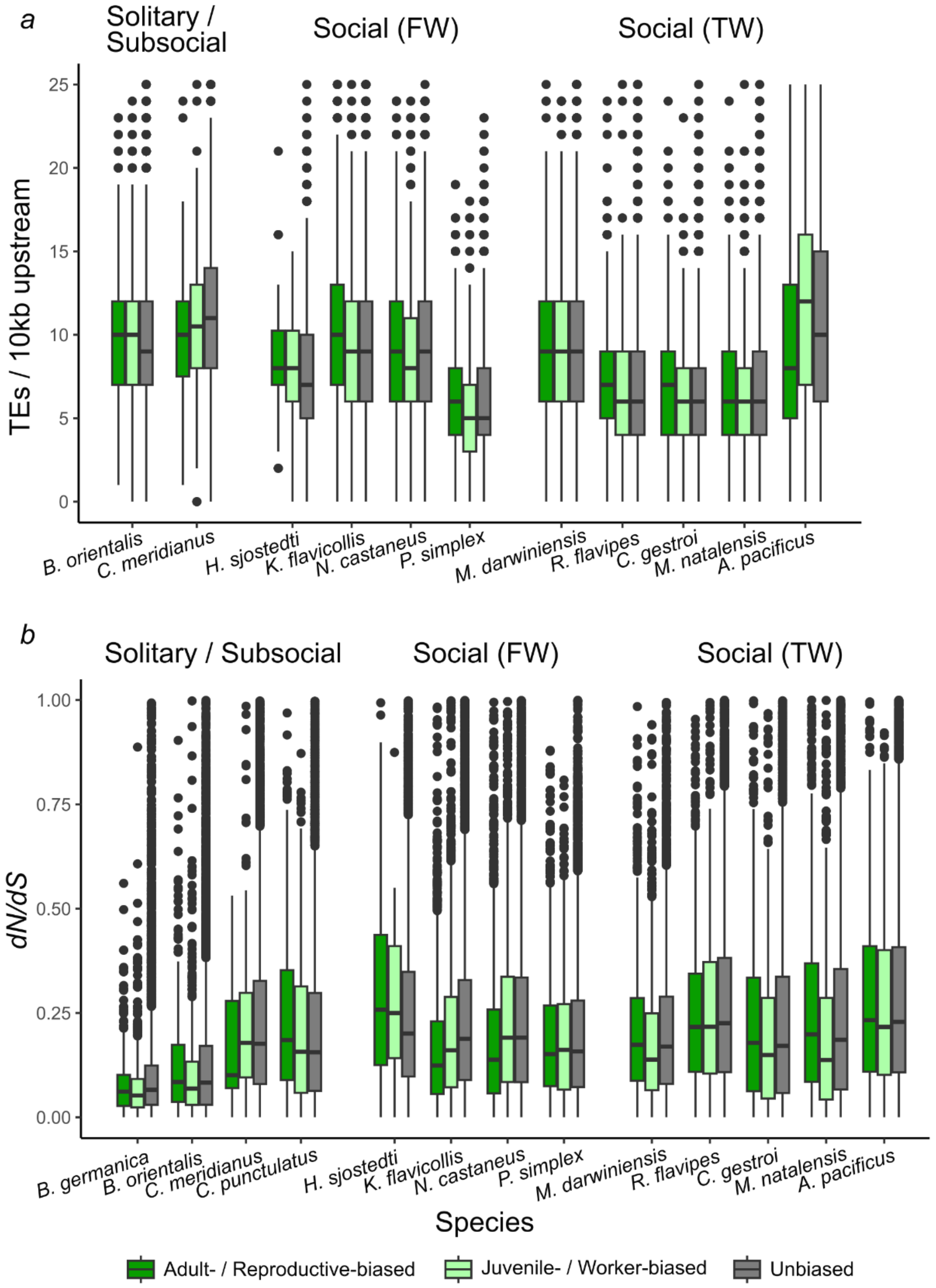

**Table S1.** Statistics of the MCMCglmm models of dependent variables *k* and *dN/dS* values from SCOs and all other HOGs. Dependent variables were tested against social category (solitary, subsocial, social (Linear, FW) and social (Bifurcated, TW)), with 3 types of priors (expanded, flat and Invert Wishart). Effective length and autocorrelation were checked to be above 200 to validate convergence of the model. Models were run in turn, changing the order of the social category level to estimate comparisons. Significant results are highlighted in bold.

| **SCO - *dN/dS* (expanded prior and with non-phylogenetic random effect)** | | | | | |
|---|---|---|---|---|---|
| | post.mean | l-95% CI | u-95% CI | eff.samp | pMCMC |
| Solitary (Intercept) | 2.5056695 | 2.083398 | 2.912499 | 2000 | 5.00E-04 |
| Subsocial | -0.5651878 | -1.27025 | 0.147814 | 2185.032 | 0.107 |
| LinearDev | -0.4839535 | -1.10433 | 0.098002 | 2147.938 | 0.102 |
| **BifurDev** | **-0.6511554** | **-1.30861** | **-0.09736** | 2121.364 | 0.036 |
| | post.mean | l-95% CI | u-95% CI | eff.samp | pMCMC |
| Subsocial (Intercept) | 1.9537106 | 1.258481 | 2.56346 | 1853.728 | 5.00E-04 |
| Solitary | 0.5545583 | -0.18991 | 1.294016 | 2000 | 0.118 |
| LinearDev | 0.0740312 | -0.50995 | 0.697046 | 2343.615 | 0.816 |
| BifurDev | -0.0989477 | -0.71335 | 0.591903 | 2398.159 | 0.719 |
| | post.mean | l-95% CI | u-95% CI | eff.samp | pMCMC |
| LinearDev (Intercept) | 2.0218427 | 1.504338 | 2.537341 | 2334.004 | 5.00E-04 |
| Solitary | 0.47725 | -0.17521 | 1.043502 | 2379.502 | 0.119 |
| Subsocial | -0.0770719 | -0.71593 | 0.558588 | 1839.601 | 0.805 |
| BifurDev | -0.1644024 | -0.45558 | 0.13602 | 2000 | 0.259 |
| | post.mean | l-95% CI | u-95% CI | eff.samp | pMCMC |
| BifurDev (Intercept) | 1.8616804 | 1.310138 | 2.35558 | 2367.732 | 5.00E-04 |
| LinearDev | 0.1628771 | -0.12038 | 0.459662 | 2000 | 0.269 |
| Solitary | 0.636326 | -0.0074 | 1.231374 | 2000 | 0.044 |
| Subsocial | 0.0879202 | -0.48175 | 0.706784 | 1857.52 | 0.756 |
| **Credible interval range calculation** | | | | | |
| Solitary (Intercept) | 0 | 0.422272 | 0.406829 | | |
| Subsocial | -0.5651878 | 0.705059 | 0.713002 | | |
| LinearDev | -0.4839535 | 0.620377 | 0.581956 | | |
| BifurDev | -0.6511554 | 0.657456 | 0.553795 | | |

| **All other HOGs – *dN/dS* (expanded prior and with non-phylogenetic random effect)** | | | | | |
|---|---|---|---|---|---|
| | post.mean | l-95% CI | u-95% CI | eff.samp | pMCMC |

| | | | | | |
|---|---|---|---|---|---|
| Solitary (Intercept) | 2.12035173 | 1.837209 | 2.4478 | 1847.465 | 5.00E-04 |
| Subsocial | -0.5168617 | -1.10752 | 0.001463 | 3078.189 | 0.063 |
| LinearDev | -0.4149854 | -0.90296 | 0.020906 | 2000 | 0.08 |
| **BifurDev** | **-0.5625463** | **-1.01759** | **-0.10028** | 2000 | 0.027 |
| | post.mean | l-95% CI | u-95% CI | eff.samp | pMCMC |
| Subsocial (Intercept) | 1.60539123 | 1.114829 | 2.105049 | 1783.528 | 5.00E-04 |
| Solitary | 0.51971964 | -0.03162 | 1.083565 | 2000 | 0.068 |
| LinearDev | 0.09213122 | -0.36827 | 0.577869 | 2000 | 0.667 |
| BifurDev | -0.0477712 | -0.52523 | 0.445477 | 2000 | 0.795 |
| | post.mean | l-95% CI | u-95% CI | eff.samp | pMCMC |
| LinearDev (Intercept) | 1.71197273 | 1.323215 | 2.11806 | 2352.572 | 5.00E-04 |
| Solitary | 0.41594252 | -0.09897 | 0.864685 | 2185.174 | 0.096 |
| Subsocial | -0.1046833 | -0.66311 | 0.333024 | 2000 | 0.664 |
| BifurDev | -0.1438966 | -0.40346 | 0.09594 | 2000 | 0.236 |
| | post.mean | l-95% CI | u-95% CI | eff.samp | pMCMC |
| BifurDev (Intercept) | 1.56468744 | 1.203661 | 1.993852 | 2167 | 5.00E-04 |
| LinearDev | 0.14379714 | -0.08771 | 0.370273 | 2080.414 | 0.214 |
| **Solitary** | **0.55960236** | **0.059168** | **1.03809** | 2000 | 0.038 |
| Subsocial | 0.04083195 | -0.43928 | 0.555279 | 2068.997 | 0.827 |
| **Credible interval range calculation** | | | | | |
| Solitary (Intercept) | 0 | 0.283143 | 0.327448 | | |
| Subsocial | -0.5168617 | 0.590661 | 0.518325 | | |
| LinearDev | -0.4149854 | 0.487979 | 0.435892 | | |
| BifurDev | -0.5625463 | 0.455048 | 0.462268 | | |

| **SCO - *k* (expanded prior and with non-phylogenetic random effect)** | | | | | |
|---|---|---|---|---|---|
| | post.mean | l-95% CI | u-95% CI | eff.samp | pMCMC |
| Solitary (Intercept) | -0.9460891 | -1.29624 | -0.67044 | 2000 | 5.00E-04 |
| **Subsocial** | **0.58832841** | **0.082724** | **1.117036** | 1736.001 | 0.031 |
| **LinearDev** | **0.77532864** | **0.302636** | **1.165372** | 2139.162 | 0.002 |
| **BifurDev** | **0.76180599** | **0.360878** | **1.189868** | 2000 | 0.004 |
| | post.mean | l-95% CI | u-95% CI | eff.samp | pMCMC |
| Subsocial (Intercept) | -0.3601537 | -0.80124 | 0.128957 | 1956.2 | 0.105 |
| **Solitary** | **-0.5870272** | **-1.16702** | **-0.11757** | 2000 | 0.034 |
| LinearDev | 0.18045782 | -0.23474 | 0.694943 | 2000 | 0.393 |
| BifurDev | 0.17372989 | -0.23921 | 0.676149 | 2000 | 0.407 |

| | | | | | |
|---|---|---|---|---|---|
| | post.mean | l-95% CI | u-95% CI | eff.samp | pMCMC |
| LinearDev (Intercept) | -0.1786203 | -0.52174 | 0.180546 | 2000 | 0.237 |
| **Solitary** | **-0.7685796** | **-1.16843** | **-0.3299** | 2000 | 0.004 |
| Subsocial | -0.1810941 | -0.68882 | 0.214925 | 1873.995 | 0.391 |
| BifurDev | -0.0057526 | -0.22272 | 0.262115 | 2000 | 0.918 |
| | post.mean | l-95% CI | u-95% CI | eff.samp | pMCMC |
| BifurDev (Intercept) | -0.1848542 | -0.53048 | 0.187614 | 2000 | 0.237 |
| LinearDev | 0.01264549 | -0.22396 | 0.249081 | 2327.589 | 0.866 |
| **Solitary** | **-0.7578418** | **-1.1836** | **-0.29156** | 2000 | 0.002 |
| Subsocial | -0.1717006 | -0.65348 | 0.289634 | 2000 | 0.395 |
| **Credible interval range calculation** | | | | | |
| Solitary (Intercept) | 0 | 0.350155 | 0.275648 | | |
| Subsocial | 0.58832841 | 0.505604 | 0.528708 | | |
| LinearDev | 0.77532864 | 0.472693 | 0.390044 | | |
| BifurDev | 0.76180599 | 0.400928 | 0.428062 | | |

**Table S2.** Test outcomes from the 569 families that were significantly evolving in the CAFE analysis. Gene counts were tested against ontogeny (binomial variable Linear or Bifurcated) and social category (Categorical variable Solitary, Subsocial, Social (FW), Social (TW)). For the latter, the p-value of the "Social (TW)" category was considered. Two types of priors were used (expanded (exp) and Invert-Wishart (iw)) with and without non-phylogenetic random factor. p-values were adjusted with BH correction. Families were only considered if *Mastotermes darwiniensis* (Mdar) and *Dolichorhinotermes longilabius* (Dlon) clustered more closely to bifurcated (TW) versus linear (FW) species (Fig. S3).

| **Covariate** | **Prior type** | **Random factor** | **HOG ID** | **p-value** | **Adjusted p-value** | **Mdar and Dlon** |
|---|---|---|---|---|---|---|
| Ontogeny | iw | with | **N0.HOG0000380** | 5.00E-04 | 0.014225 | Yes |
| Ontogeny | iw | with | N0.HOG0000481 | 5.00E-04 | 0.014225 | No |
| Ontogeny | iw | with | N0.HOG0000502 | 5.00E-04 | 0.014225 | No |
| Ontogeny | iw | with | N0.HOG0000659 | 5.00E-04 | 0.014225 | No |
| Ontogeny | iw | with | N0.HOG0000906 | 0.002 | 0.0355625 | No |
| Ontogeny | iw | with | **N0.HOG0000937** | 0.001 | 0.02276 | Yes |
| Ontogeny | iw | with | N0.HOG0000964 | 0.001 | 0.02276 | No |
| Ontogeny | iw | with | N0.HOG0000974 | 5.00E-04 | 0.014225 | No |
| Ontogeny | iw | with | N0.HOG0000976 | 0.001 | 0.02276 | No |
| Ontogeny | iw | with | N0.HOG0001119 | 0.003 | 0.044921053 | No |
| Ontogeny | iw | with | N0.HOG0001245 | 5.00E-04 | 0.014225 | No |
| Ontogeny | iw | with | N0.HOG0001248 | 0.001 | 0.02276 | No |
| Ontogeny | iw | with | N0.HOG0001292 | 5.00E-04 | 0.014225 | No |
| Ontogeny | iw | with | **N0.HOG0001513** | 0.002 | 0.0355625 | Yes |
| Ontogeny | iw | with | N0.HOG0001526 | 0.002 | 0.0355625 | No |
| Ontogeny | iw | with | N0.HOG0001537 | 0.002 | 0.0355625 | No |
| Ontogeny | iw | with | N0.HOG0001575 | 0.003 | 0.044921053 | No |
| Ontogeny | iw | with | N0.HOG0001671 | 5.00E-04 | 0.014225 | No |
| Ontogeny | iw | with | N0.HOG0001738 | 5.00E-04 | 0.014225 | No |
| Ontogeny | iw | with | N0.HOG0001798 | 5.00E-04 | 0.014225 | No |
| Ontogeny | iw | with | N0.HOG0001834 | 5.00E-04 | 0.014225 | No |
| Ontogeny | iw | with | **N0.HOG0001851** | 5.00E-04 | 0.014225 | Yes |
| Ontogeny | iw | with | N0.HOG0001878 | 5.00E-04 | 0.014225 | No |
| Ontogeny | iw | with | N0.HOG0001979 | 5.00E-04 | 0.014225 | No |
| Ontogeny | iw | with | N0.HOG0002048 | 5.00E-04 | 0.014225 | No |
| Ontogeny | iw | with | N0.HOG0002107 | 5.00E-04 | 0.014225 | No |
| Ontogeny | iw | with | N0.HOG0002323 | 0.002 | 0.0355625 | No |
| Ontogeny | iw | with | N0.HOG0002350 | 0.003 | 0.044921053 | No |
| Ontogeny | iw | with | N0.HOG0002353 | 5.00E-04 | 0.014225 | No |
| Ontogeny | iw | with | N0.HOG0002845 | 5.00E-04 | 0.014225 | No |
| Ontogeny | iw | with | N0.HOG0002865 | 0.002 | 0.0355625 | No |

| | | | | | | |
|---|---|---|---|---|---|---|
| Ontogeny | iw | with | **N0.HOG0002943** | 0.001 | 0.02276 | Yes |
| Ontogeny | iw | with | N0.HOG0003014 | 0.002 | 0.0355625 | No |
| Ontogeny | iw | with | N0.HOG0003077 | 5.00E-04 | 0.014225 | No |
| Ontogeny | iw | with | N0.HOG0003085 | 0.003 | 0.044921053 | No |
| Ontogeny | iw | with | N0.HOG0003410 | 5.00E-04 | 0.014225 | No |
| Ontogeny | iw | with | **N0.HOG0004986** | 0.003 | 0.044921053 | Yes |
| Ontogeny | iw | with | **N0.HOG0006820** | 0.003 | 0.044921053 | Yes |
| Ontogeny | iw | without | N0.HOG0000225 | 0.001 | 0.018966667 | No |
| Ontogeny | iw | without | **N0.HOG0000380** | 5.00E-04 | 0.013547619 | Yes |
| Ontogeny | iw | without | N0.HOG0000481 | 5.00E-04 | 0.013547619 | No |
| Ontogeny | iw | without | N0.HOG0000502 | 5.00E-04 | 0.013547619 | No |
| Ontogeny | iw | without | N0.HOG0000659 | 5.00E-04 | 0.013547619 | No |
| Ontogeny | iw | without | **N0.HOG0000758** | 0.003 | 0.044921053 | Yes |
| Ontogeny | iw | without | N0.HOG0000781 | 0.003 | 0.044921053 | No |
| Ontogeny | iw | without | N0.HOG0000906 | 0.002 | 0.033470588 | No |
| Ontogeny | iw | without | **N0.HOG0000937** | 0.001 | 0.018966667 | Yes |
| Ontogeny | iw | without | N0.HOG0000964 | 5.00E-04 | 0.013547619 | No |
| Ontogeny | iw | without | N0.HOG0000974 | 0.001 | 0.018966667 | No |
| Ontogeny | iw | without | N0.HOG0000976 | 5.00E-04 | 0.013547619 | No |
| Ontogeny | iw | without | N0.HOG0001119 | 0.001 | 0.018966667 | No |
| Ontogeny | iw | without | N0.HOG0001245 | 0.001 | 0.018966667 | No |
| Ontogeny | iw | without | N0.HOG0001248 | 5.00E-04 | 0.013547619 | No |
| Ontogeny | iw | without | N0.HOG0001292 | 5.00E-04 | 0.013547619 | No |
| Ontogeny | iw | without | N0.HOG0001476 | 0.003 | 0.044921053 | No |
| Ontogeny | iw | without | **N0.HOG0001513** | 0.001 | 0.018966667 | Yes |
| Ontogeny | iw | without | N0.HOG0001526 | 5.00E-04 | 0.013547619 | No |
| Ontogeny | iw | without | N0.HOG0001575 | 0.001 | 0.018966667 | No |
| Ontogeny | iw | without | N0.HOG0001671 | 5.00E-04 | 0.013547619 | No |
| Ontogeny | iw | without | N0.HOG0001738 | 5.00E-04 | 0.013547619 | No |
| Ontogeny | iw | without | N0.HOG0001798 | 5.00E-04 | 0.013547619 | No |
| Ontogeny | iw | without | N0.HOG0001834 | 0.001 | 0.018966667 | No |
| Ontogeny | iw | without | **N0.HOG0001851** | 5.00E-04 | 0.013547619 | Yes |
| Ontogeny | iw | without | N0.HOG0001878 | 5.00E-04 | 0.013547619 | No |
| Ontogeny | iw | without | N0.HOG0001979 | 0.001 | 0.018966667 | No |
| Ontogeny | iw | without | N0.HOG0002048 | 5.00E-04 | 0.013547619 | No |
| Ontogeny | iw | without | N0.HOG0002107 | 5.00E-04 | 0.013547619 | No |
| Ontogeny | iw | without | N0.HOG0002350 | 0.002 | 0.033470588 | No |
| Ontogeny | iw | without | N0.HOG0002353 | 5.00E-04 | 0.013547619 | No |
| Ontogeny | iw | without | N0.HOG0002845 | 5.00E-04 | 0.013547619 | No |
| Ontogeny | iw | without | N0.HOG0002865 | 5.00E-04 | 0.013547619 | No |
| Ontogeny | iw | without | **N0.HOG0002943** | 0.002 | 0.033470588 | Yes |

| | | | | | | |
|---|---|---|---|---|---|---|
| Ontogeny | iw | without | N0.HOG0003077 | 5.00E-04 | 0.013547619 | No |
| Ontogeny | iw | without | N0.HOG0003085 | 0.002 | 0.033470588 | No |
| Ontogeny | iw | without | N0.HOG0003410 | 5.00E-04 | 0.013547619 | No |
| Ontogeny | iw | without | **N0.HOG0006820** | 0.003 | 0.044921053 | Yes |
| Social_category | iw | with | **N0.HOG0000380** | 5.00E-04 | 0.0284 | Yes |
| Social_category | iw | with | N0.HOG0000502 | 5.00E-04 | 0.0284 | No |
| Social_category | iw | with | N0.HOG0000976 | 0.001 | 0.037866667 | No |
| Social_category | iw | with | N0.HOG0001248 | 0.001 | 0.037866667 | No |
| Social_category | iw | with | N0.HOG0001292 | 5.00E-04 | 0.0284 | No |
| Social_category | iw | with | N0.HOG0001671 | 5.00E-04 | 0.0284 | No |
| Social_category | iw | with | N0.HOG0001738 | 0.001 | 0.037866667 | No |
| Social_category | iw | with | N0.HOG0001798 | 5.00E-04 | 0.0284 | No |
| Social_category | iw | with | **N0.HOG0001851** | 5.00E-04 | 0.0284 | Yes |
| Social_category | iw | with | **N0.HOG0002046** | 0.001 | 0.037866667 | Yes |
| Social_category | iw | with | N0.HOG0002353 | 5.00E-04 | 0.0284 | No |
| Social_category | iw | with | **N0.HOG0002649** | 5.00E-04 | 0.0284 | Yes |
| Social_category | iw | with | N0.HOG0003077 | 5.00E-04 | 0.0284 | No |
| Social_category | iw | with | N0.HOG0003410 | 5.00E-04 | 0.0284 | No |
| Social_category | iw | with | **N0.HOG0004986** | 0.001 | 0.037866667 | Yes |
| Social_category | iw | without | **N0.HOG0000380** | 5.00E-04 | 0.0355625 | Yes |
| Social_category | iw | without | N0.HOG0000481 | 0.001 | 0.0355625 | No |
| Social_category | iw | without | N0.HOG0000502 | 5.00E-04 | 0.0355625 | No |
| Social_category | iw | without | N0.HOG0000937 | 0.001 | 0.0355625 | No |
| Social_category | iw | without | N0.HOG0000976 | 0.001 | 0.0355625 | No |
| Social_category | iw | without | N0.HOG0001248 | 0.001 | 0.0355625 | No |
| Social_category | iw | without | N0.HOG0001292 | 0.001 | 0.0355625 | No |
| Social_category | iw | without | N0.HOG0001738 | 0.001 | 0.0355625 | No |
| Social_category | iw | without | N0.HOG0001798 | 5.00E-04 | 0.0355625 | No |
| Social_category | iw | without | **N0.HOG0001851** | 5.00E-04 | 0.0355625 | Yes |
| Social_category | iw | without | **N0.HOG0002046** | 5.00E-04 | 0.0355625 | Yes |
| Social_category | iw | without | N0.HOG0002107 | 0.001 | 0.0355625 | No |
| Social_category | iw | without | N0.HOG0002353 | 5.00E-04 | 0.0355625 | No |
| Social_category | iw | without | **N0.HOG0002649** | 5.00E-04 | 0.0355625 | Yes |
| Social_category | iw | without | N0.HOG0003077 | 0.001 | 0.0355625 | No |
| Social_category | iw | without | N0.HOG0003410 | 5.00E-04 | 0.0355625 | No |

**Table S3.** Results of the significant tests from the OR families that were significantly evolving in the CAFE analysis. Gene counts were tested against ontogeny and social category. For the latter, the p-value of the bifurcated category was considered. Two types of priors were used (expanded and Invert Wishart) with and without a non-phylogenetic random factor. p-values were adjusted using a BH correction and each test was inspected to ensure that *Mastotermes darwiniensis* (Mdar) and *Dolichorhinotermes longilabius* (Dlon) clustered more closely to bifurcated (TW) versus linear (FW) species.

| **Covariate** | **Prior type** | **Random factor** | **OG ID** | **p-value** | **Adjusted p-value** | **Mdar and Dlon** |
|---|---|---|---|---|---|---|
| Ontogeny | iw | with | Gene Cluster 2 | 5.00E-04 | 0.021667 | No |
| Ontogeny | iw | with | Gene Cluster 1 (OG112) | 0.001 | 0.021667 | Yes |
| Ontogeny | iw | with | OG202 | 0.001 | 0.021667 | No |
| Ontogeny | iw | with | OG21 | 0.003 | 0.04875 | Yes |
| Ontogeny | iw | without | Gene Cluster 2 | 5.00E-04 | 0.010833 | No |
| Ontogeny | iw | without | Gene Cluster 1 (OG112) | 5.00E-04 | 0.010833 | Yes |
| Ontogeny | iw | without | OG202 | 5.00E-04 | 0.010833 | No |
| Ontogeny | iw | without | OG21 | 0.001 | 0.01625 | Yes |
| Social_category | iw | with | Gene Cluster 2 | 5.00E-04 | 0.01625 | No |
| Social_category | iw | with | Gene Cluster 1 (OG112) | 5.00E-04 | 0.01625 | Yes |
| Social_category | iw | without | Gene Cluster 2 | 5.00E-04 | 0.01625 | No |
| Social_category | iw | without | Gene Cluster 1 (OG112) | 5.00E-04 | 0.01625 | Yes |

**Table S4.** Selection analysis results of OR-OGs. p-value indicates the result of aBSREL test for positive selection per OR cluster.

| **OR-OG** | **Species** | **Gene ID** | **p-value** |
|---|---|---|---|
| OR-cluster 1 (OR-OG112) | *Mastotermes darwiniensis* | MdarORsg142_t1 | 0 |
| OR-cluster 1 (OR-OG112) | *Mastotermes darwiniensis* | Mdar00005033s1_R0_split1 | 0.00006 |
| OR-cluster 1 (OR-OG112) | *Dolichorhinotermes longilabius* | Dlon00014506_R0 | 0.00018 |
| OR-cluster 1 (OR-OG112) | *Macrotermes natalensis* | Mnat00004924_R0 | 0.00022 |
| OR-cluster 1 (OR-OG112) | *Sphaerotermes sphaerothorax* | Ssph00007962_R0 | 0.00085 |
| OR-cluster 1 (OR-OG112) | *Coptotermes testaceus* | Ctes00008990_R0 | 0.01288 |
| OR-cluster 1 (OR-OG112) | *Mastotermes darwiniensis* | Mdar00005022_R0 | 0.03707 |
| OR-cluster 2 | *Anoplotermes pacificus* | Apac00000430_R0 | 0.00426 |
| OR-cluster 2 | *Coptotermes gestroi* | Cges00000741_R0 | 0.00523 |
| OR-cluster 2 | *Macrotermes natalensis* | Mnat00000840_R0 | 0.03878 |
| OR-OG202 | *Reticulitermes flavipes* | RflaORsg126_t1 | 0 |
| OR-OG202 | *Anoplotermes pacificus* | ApacORsg114_t1 | 0.00023 |
| OR-OG202 | *Neocapritermes taracua* | Ntar00020396_R0 | 0.03113 |
| OR-OG202 | *Macrotermes natalensis* | Mnat00000840_R0 | 0.0417 |
| OR-OG21 | *Reticulitermes flavipes* | Rfla00001431_R0 | 0 |
| OR-OG21 | *Coatitermes sp.* | Csp400001497_R0 | 0 |
| OR-OG21 | *Hodotermopsis sjostedti* | Hsjo00015839_R0 | 0.00645 |

**Table. S5.** Connectivity between worker- and reproductive-specific ORs and DEGs in pairwise comparisons of worker and reproductive networks ("Network caste"). Worker or reproductive-specific ORs ("OR caste") as well as worker- or reproductive-biased DEGs shared in reproductives or workers of the social category of a given species (i.e. FW or TW) were subset from each network. Connectivity between DEGs that were either shared between reproductives or workers (DEG caste) and either total ORs, OR-cluster 1 and 2, was calculated, and divided by the number of ORs and then by the number of DEGs, yielding degrees of connectivity for total ORs ("Connectivity - All ORs), OR-cluster 1 ("Connectivity -  Cluster 1 (OG-OR112)") and OR-cluster 2 ("Connectivity - Cluster 2"), respectively.

| Network caste | OR caste | DEG caste | Species | Connectivity - All ORs | Connectivity - OR-cluster 1 | Connectivity - Cluster 2 |
|---|---|---|---|---|---|---|
| Wo | Wo | Wo | Apac | 0.74 | 0.75 | 0.76 |
| Re | Re | Re | Apac | 0.28 | 0,53 | 0,29 |
| Wo | Wo | Wo | Cges | 0,74 | 0,68 | 0,66 |
| Re | Re | Re | Cges | 0,58 | 0,52 | 0,58 |
| Wo | Wo | Wo | Hsjo | 0,27 | 0,28 | 0 |
| Re | Re | Re | Hsjo | 0,19 | 0,15 | 0 |
| Wo | Wo | Wo | Kfla | 0,23 | 0,22 | 0 |
| Re | Re | Re | Kfla | 0,24 | 0,08 | 0 |
| Wo | Wo | Wo | Mdar | 0,22 | 0,19 | 0 |
| Re | Re | Re | Mdar | 0,17 | 0,18 | 0 |
| Wo | Wo | Wo | Mnat | 0,41 | 0,36 | 0,32 |
| Re | Re | Re | Mnat | 0,53 | 0,61 | 0,49 |
| Wo | Wo | Wo | Ncas | 0,3 | 0 | 0 |
| Re | Re | Re | Ncas | 0,37 | 0,31 | 0 |
| Wo | Wo | Wo | PRsim | 0.22 | 0.19 | 0.28 |
| Re | Re | Re | PRsim | 0.45 | 0.43 | 0.33 |
| Wo | Wo | Wo | Rfla | 0.65 | 0.43 | 0.77 |
| Re | Re | Re | Rfla | 0.27 | 0.31 | 0.28 |

**Table S6.** Generalized linear mixed model analysis of *dN/dS* and TE overlap. Pairwise comparisons by social category (solitary cockroaches, subsocial wood roaches, linear (FW) and bifurcated (TW) termites) and gene expression bias (Reproductive vs. Worker vs. Unbiased and Adult vs. Juvenile vs. Unbiased in cockroaches, respectively). For TEs, solitary cockroaches and subsocial wood roaches were combined into a single category. Significant comparisons are indicated in bold.

| | **Expression bias/Development group** | **Contrast** | **Estimate** | **SE** | **Z ratio** | **P-value** |
|---|---|---|---|---|---|---|
| dN/dS | Unbiased | Linear – Bifur | -0.030852 | 0.0274 | -1.128 | 0.6723 |
| | | **Linear – Cockroaches** | **0.106097** | 0.0353 | 3.004 | **0.0142** |
| | | Linear – wood roaches | -0.002672 | 0.0353 | -0.076 | 0.9998 |
| | | **Bifur – Cockroaches** | **0.136949** | **0.0341** | **4.014** | **0.0003** |
| | | Bifur – wood roaches | 0.028180 | 0.0341 | 0.827 | 0.8418 |
| | | **Roaches – wood roaches** | **-0.108769** | **0.0408** | **-2.669** | **0.0381** |
| | Reproductive | **Linear – Bifur** | **-0.097972** | **0.0285** | **-3.441** | **0.0032** |
| | | Linear – Cockroaches | 0.072573 | 0.0384 | 1.892 | 0.2315 |
| | | **Linear - wood roaches** | **-0.124193** | **0.0405** | **-3.065** | **0.0117** |
| | | **Bifur – Cockroaches** | **0.170544** | **0.0373** | **4.575** | **<.0001** |
| | | Bifur – wood roaches | -0.026221 | 0.0395 | -0.664 | 0.9106 |
| | | **Roaches - wood roaches** | **-0.196766** | **0.0471** | **-4.175** | **0.0002** |
| | Worker | Linear – Bifur | -0.005100 | 0.0286 | -0.178 | 0.9980 |
| | | **Linear – Cockroaches** | **0.148902** | **0.0375** | **3.975** | **0.0004** |
| | | Linear – wood roaches | -0.000928 | 0.0419 | -0.022 | 1.0000 |
| | | **Bifur – Cockroaches** | **0.154001** | **0.0363** | **4.242** | **0.0001** |
| | | Bifur – wood roaches | 0.004172 | 0.0408 | 0.102 | 0.9996 |
| | | **Cockroaches – wood roaches** | -0.149829 | 0.0474 | -3.159 | 0.0086 |
| | Termites with linear development | **Unbiased – Repr** | **0.0787** | **0.00656** | **11.998** | **<.0001** |
| | | **Unbiased – Wo** | **0.0252** | **0.00706** | **3.576** | **0.0010** |
| | | **Repr – Wo** | **-0.0535** | **0.00865** | **-6.184** | **<.0001** |
| | Termites with bifurcated development | Unbiased – Repr | 0.0116 | 0.00640 | 1.816 | 0.1644 |
| | | **Unbiased – Wo** | **0.0510** | **0.00668** | **7.637** | **<.0001** |

| | | | | | | |
|---|---|---|---|---|---|---|
| | | **Repr – Wo** | **0.0394** | **0.00868** | **4.538** | **<.0001** |
| | Cockroaches | **Unbiased – Adult** | **0.0452** | **0.01470** | **3.068** | **0.0061** |
| | | **Unbiased – Juvenile** | **0.0680** | **0.01190** | **5.723** | **<.0001** |
| | | Adult – Juvenile | 0.0228 | 0.01830 | 1.247 | 0.4255 |
| | wood roaches | Unbiased - Adult | -0.0428 | 0.01960 | -2.183 | 0.0741 |
| | | Unbiased - Juvenile | 0.0270 | 0.02210 | 1.221 | 0.4405 |
| | | **Adult – Juvenile** | **0.0698** | **0.02920** | **2.391** | **0.0443** |
| TE | Unbiased | Linear – Bifur | -0.126 | 1.20 | -0.105 | 0.9940 |
| | | Linear – Roaches | -2.179 | 1.55 | -1.404 | 0.3387 |
| | | Bifur – Roaches | -2.053 | 1.50 | -1.370 | 0.3569 |
| | Reproductive | Linear – Bifur | 0.450 | 1.20 | 0.374 | 0.9260 |
| | | Linear – Roaches | -2.314 | 1.56 | -1.481 | 0.3002 |
| | | Bifur – Roaches | -2.764 | 1.51 | -1.829 | 0.1600 |
| | Worker | Linear – Bifur | -0.304 | 1.20 | -0.252 | 0.9656 |
| | | Linear – Roaches | -2.807 | 1.56 | -1.802 | 0.1690 |
| | | Bifur – Roaches | -2.504 | 1.51 | -1.663 | 0.2196 |
| | Termites with linear development | **Unbiased – Repr** | **-0.4952** | **0.0587** | **-8.433** | **<.0001** |
| | | **Unbiased – Wo** | **0.1457** | **0.0618** | **2.356** | **0.0484** |
| | | **Repr – Wo** | **0.6409** | **0.0782** | **8.200** | **<.0001** |
| | Termites with bifurcated development | Unbiased – Repr | 0.0803 | 0.0582 | 1.380 | 0.3517 |
| | | Unbiased – Wo | -0.0321 | 0.0601 | -0.534 | 0.8546 |
| | | Repr – Wo | -0.1124 | 0.0797 | -1.410 | 0.3358 |
| | Cockroaches | **Unbiased – Adult** | **-0.6304** | **0.1820** | **-3.457** | **0.0016** |
| | | **Unbiased – Juvenile** | **-0.4827** | **0.1330** | **-3.617** | **0.0009** |
| | | Adult – Juvenile | 0.1477 | 0.2240 | 0.660 | 0.7865 |

**Table S7.** Number and identity of HOGs shared between i) all TW species and their representation in corresponding FW species, ii) all FW species and their representation in corresponding TW species for worker- (top two panels) and reproductive-biased (bottom two panels) DEGs.

| Worker-biased HOG in all TW species | Number of FW species the HOG is shared with | Name of the species the HOG is shared with |
|---|---|---|
| HOG0003327 | 1 | Ncas |
| HOG0005566 | 0 | |
| HOG0009626 | 1 | PRsim |
| HOG0012340 | 1 | Kfla |
| HOG0013615 | 1 | PRsim |
| HOG0014907 | 2 | Ncas, Kfla |
| HOG0015385 | 1 | Kfla |
| mean | *1* | |
| Worker-biased HOG in all FW species | Number of TW species the HOG is shared with | Name of the species the HOG is shared with |
| HOG0000495 | 0 | |
| HOG0001361 | 4 | Rfla, Mnat, Cges, Apac |
| HOG0009391 | 4 | Rfla, Mnat, Mdar, Cges |
| HOG0011646 | 2 | Mdar, Apac |
| HOG0013829 | 4 | Mnat, Mdar, Cges, Apac |
| mean | *2.8* | |
| Reproductive-biased HOG in all TW species | Number of FW species the HOG is shared with | Name of the species the HOG is shared with |
| HOG0003220 | 3 | PRsim, Kfla, Hsjo |
| HOG0003766 | 2 | Ncas, Kfla |
| HOG0003789 | 4 | PRsim, Ncas, Kfla, Hsjo |
| HOG0006583 | 1 | PRsim |
| HOG0008018 | 0 | |
| HOG0008558 | 2 | Ncas, Kfla |
| HOG0008956 | 0 | |
| HOG0010194 | 3 | PRsim, Ncas, Kfla |
| HOG0012362 | 1 | Ncas |
| HOG0012788 | 0 | |
| HOG0014377 | 3 | PRsim, Ncas, Kfla |
| HOG0014765 | 4 | PRsim, Ncas, Kfla, Hsjo |
| HOG0015687 | 2 | PRsim, Kfla |
| HOG0015940 | 2 | PRsim, Kfla |
| HOG0016311 | 3 | PRsim, Ncas, Hsjo |
| mean | *2* | |

| | | |
|---|---|---|
| Reproductive-biased HOG in all FW species | Number of TW species the HOG is shared with | Name of the species the HOG is shared with |
| HOG0002112 | 3 | Rfla, Mdar, Apac |
| HOG0002381 | 3 | Mnat, Mdar, Apac |
| HOG0003748 | 1 | Rfla |
| HOG0003789 | 5 | Rfla, Mnat, Mdar, Cges, Apac |
| HOG0010345 | 2 | Mdar, Apac |
| HOG0012170 | 4 | Rfla, Mnat, Mdar, Cges |
| HOG0014765 | 5 | Rfla, Mnat, Mdar, Cges, Apac |
| mean | *3.3* | |

**Table S8.** List of fossils used to calibrate the phylogeny used in this study.

| | Species | Minimum age constraint (MY) | Calibration group | Soft maximum bound (97.5% probability) | Note on maximum bound | Reference |
|---|---|---|---|---|---|---|
| USED | *Valditermes brenanae* | 125.5 | Isoptera + *Cryptocercus* | 235 | First fossil of Mesoblattinidae (*Triassoblatta argentina:* Martins-Neto et al. 2005) | Jarzembowski 1981 |
| USED | *Melqartitermes myrrheus* | 125.5 | Mastotermitidae + sister group | 235 | First fossil of Mesoblattinidae (*Triassoblatta argentina:* Martins-Neto et al. 2005) | Engel et al. 2007 |
| USED | *Cosmotermes multus* | 93.5 | Teletisoptera + sister group | 235 | First fossil of Mesoblattinidae (*Triassoblatta argentina:* Martins-Neto et al. 2005) | Zhao et al. 2019 |
| USED | *Archeorhinotermes rossi* | 94.3 | Kalotermitidae + sister group | 125.5 | First termite fossil | Krishna and Grimaldi 2003 |
| USED | *Nanotermes isaacae* | 47.8 | Termitidae + *Coptotermes + Heterotermes + Reticulitermes* | 94.3 | First fossil of Rhinotermitinae | Engel et al. 2011 |
| USED | *Huguenotermes septimaniensis* | 33.9 | *Cryptotermes* + sister group | 94.3 | First fossil of Kalotermitidae | Engel and Nel 2015 |
| USED | *Reticulitermes antiquus* | 33.9 | *Reticulitermes + Coptotermes + Heterotermes* | 94.3 | First fossil of Rhinotermitinae | Engel et al. 2007 |
| USED | *Coptotermes sucineus* | 16 | *Coptotermes + Heterotermes* | 33.9 | First fossil of *Heterotermes* | Engel 2008 |
| USED | *Constrictotermes electroconstrictus* | 13.7 | *Constrictotermes* + sister group | 47.8 | First fossil of Termitidae | Krishna 1996 |
| USED | *Anoplotermes* sensu lato | 13.7 | *Anoplotermes-*group+sister group | 47.8 | First fossil of Termitidae | Krishna and Grimaldi 2009 |
| USED | *Microcerotermes insulanus* | 13.7 | *Microcerotermes* + sister group | 47.8 | First fossil of Termitidae | Krishna and Grimaldi 2009 |
| NOT USED | *Amitermes lucidus* | 13.7 | *Amitermes +* sister group | 47.8 | First fossil of Termitidae | Krishna and Grimaldi 2009 |
| USED | *Dolichorhinotermes dominicanus* | 13.7 | *Dolichorhinotermes* + sister group | 94.3 | First fossil of Rhinotermitinae | Schlemmermeyer and Cancello 2000 |
| USED | *Macrotermes pristinus* | 11.6 | *Macrotermes* + sister group | 47.8 | First fossil of Termitidae | Charpentier 1843 |
